# Distinct tumor architectures for metastatic colonization of the brain

**DOI:** 10.1101/2023.01.27.525190

**Authors:** Siting Gan, Danilo G. Macalinao, Sayyed Hamed Shahoei, Lin Tian, Xin Jin, Harihar Basnet, James T. Muller, Pranita Atri, Evan Seffar, Walid Chatila, Anna-Katerina Hadjantonakis, Nikolaus Schultz, Edi Brogi, Tejus A. Bale, Dana Pe’er, Joan Massagué

## Abstract

Brain metastasis is a dismal cancer complication, hinging on the initial survival and outgrowth of disseminated cancer cells. To understand these crucial early stages of colonization, we investigated two prevalent sources of cerebral relapse, triple-negative (TNBC) and HER2+ breast cancer (HER2BC). We show that these tumor types colonize the brain aggressively, yet with distinct tumor architectures, stromal interfaces, and autocrine growth programs. TNBC forms perivascular sheaths with diffusive contact with astrocytes and microglia. In contrast, HER2BC forms compact spheroids prompted by autonomous extracellular matrix components and segregating stromal cells to their periphery. Single-cell transcriptomic dissection reveals canonical Alzheimer’s disease-associated microglia (DAM) responses. Differential engagement of tumor-DAM signaling through the receptor AXL suggests specific pro-metastatic functions of the tumor architecture in both TNBC perivascular and HER2BC spheroidal colonies. The distinct spatial features of these two highly efficient modes of brain colonization have relevance for leveraging the stroma to treat brain metastasis.

## Introduction

Brain metastasis is an ominous form of cancer progression, with severe neurological complications, dismal survival rates, and limited treatment options^1, 2^. It is the most common malignancy in the central nervous system (CNS)^3^, and frequently occurs in patients with breast cancer, lung cancer, and melanoma^1^. The risk of brain metastasis depends on the specific tumor type. For example, 20-30% of patients with the basal subtypes HER2+ breast cancer (HER2BC) or triple-negative breast cancer (TNBC) develop brain metastasis, whereas patients with luminal breast cancer subtypes do so infrequently (under 10% of cases), even though blood circulation patterns facilitate the dissemination of these tumors to the brain equally^4, 5^. Effective treatment options for residual breast cancer are limited. TNBC lacks sufficient expression of the estrogen receptor, progesterone receptor or the receptor HER2 for hormone therapy or drugs targeting HER2 to work^6^. In HER2BC patients, due to the restricted permeability of blood-brain barrier, systemic treatments with anti-HER2 antibody are effective against disseminated disease in visceral organs but less effective against relapse in the brain^7–9^. Therapeutic approaches based on leveraging immune components of the cerebral parenchyma could offer promise but have been hampered by a lack of knowledge about the brain metastatic stroma, particularly in the context of minimal residual disease.

Despite the prevalence and poor prognosis in patients, brain metastasis is a highly inefficient process at the cellular level, with the majority of disseminated cancer cells succumbing to physical, metabolic, or immunologic challenges^10^. Work to understand the basis for brain metastasis has focused on molecular mechanisms that allow disseminated cancer cells to overcome these barriers. Studies on mouse models and clinical tissue samples have identified molecular mediators of cancer cell interactions with the brain vasculature^11–14^, astrocytes^15–19^, microglia^20–25^, and neurons^26^, metabolic adaptation of metastatic cells in the brain^27–29^ and the cellular composition of the tumor microenvironment (TME) in large macrometastatic lesions^30–34^.

However, even with the aforementioned progress, the crucial early stages of brain metastatic colonization, when elimination is the predominant fate of disseminated cancer cells and their survival is on the balance, remain obscure^20, 22, 35^. In particular, the spatial features of brain metastatic colony formation and their role in disease progression are unknown. The tumor growth and stromal crosstalk are functionally embedded in tumor architecture, in that the spatial features of a colony can be both cause and consequence of various other determinants (e.g., immune infiltration) of tumor development^36^.

Here we focus on the early stages of brain colonization by two subtypes of breast cancer with high incidence of brain metastasis, TNBC and HER2BC. We report two strikingly different forms of brain colony architecture – perivascular versus spheroidal – that are differentially adopted by TNBC and HER2BC cells. These two colonization patterns create distinct spatial interfaces with the brain parenchyma, distinguished by infiltrative and segregated TME interfaces, respectively. Focusing on microglia as a prominent, highly reactive immune component of the TME, we show that both TNBC and HER2BC acutely yet differently activate microglial responses characteristic of Alzheimer’s disease. Our findings illuminate distinct strategies of brain colonization and microglia engagement by two major breast cancer subtypes, highlighting the importance of tumor spatial considerations in future efforts to eliminate metastatic disease in the brain.

## Results

### Tumor type-dependent perivascular and spheroidal brain colonization patterns

Upon extravasating from blood capillaries, metastasis-initiating cells from various types of carcinoma occupy perivascular niches to establish metastatic colonies, which is particularly apparent in brain metastasis^37^. The brain metastasis (BrM) models that we previously developed from human H2030-BrM lung adenocarcinoma (LUAD) and MDA-MB-231-BrM (MDA231-BrM for short) TNBC cells in athymic mice, and mouse E0771-BrM TNBC cells in immunocompetent mice (Figures 1A, 1B, S1A, S1B, and Supplementary videos 1-3) exemplified the vascular cooptive growth pattern. As previously reported by us^12, 13^ and others^20, 37–39^, extravasated cells migrate over the abluminal surface of capillaries, spread on the vascular basement membrane to initiate proliferation, and form sheaths around the vessels that engulf the local capillary network before eventually transitioning towards a multi-layered colony structure. 2D imaging of brain slices using confocal microscopy (Figure S1A) and 3D imaging of cleared whole brain hemispheres using light-sheet microscopy (Figure S1B and Supplementary videos 1-3) revealed that BrM cancer cells disseminated to the mouse brain through the blood circulation stochastically formed individual metastatic colonies spanning a range of sizes (with a radius from 10 to 1000 µm), which allowed sampling a multitude of micrometastatic colonies even at a single time point from one mouse. Spreading of the cancer cells on the perivascular basement membrane is mediated by the cell adhesion molecules L1CAM and β1-integrins binding to perivascular basement membrane laminins^12, 40^, which triggers activation of the transcription factors YAP and MRTF in metastasis-initiating cells for tumor colony outgrowth^13^.

**Figure 1.**
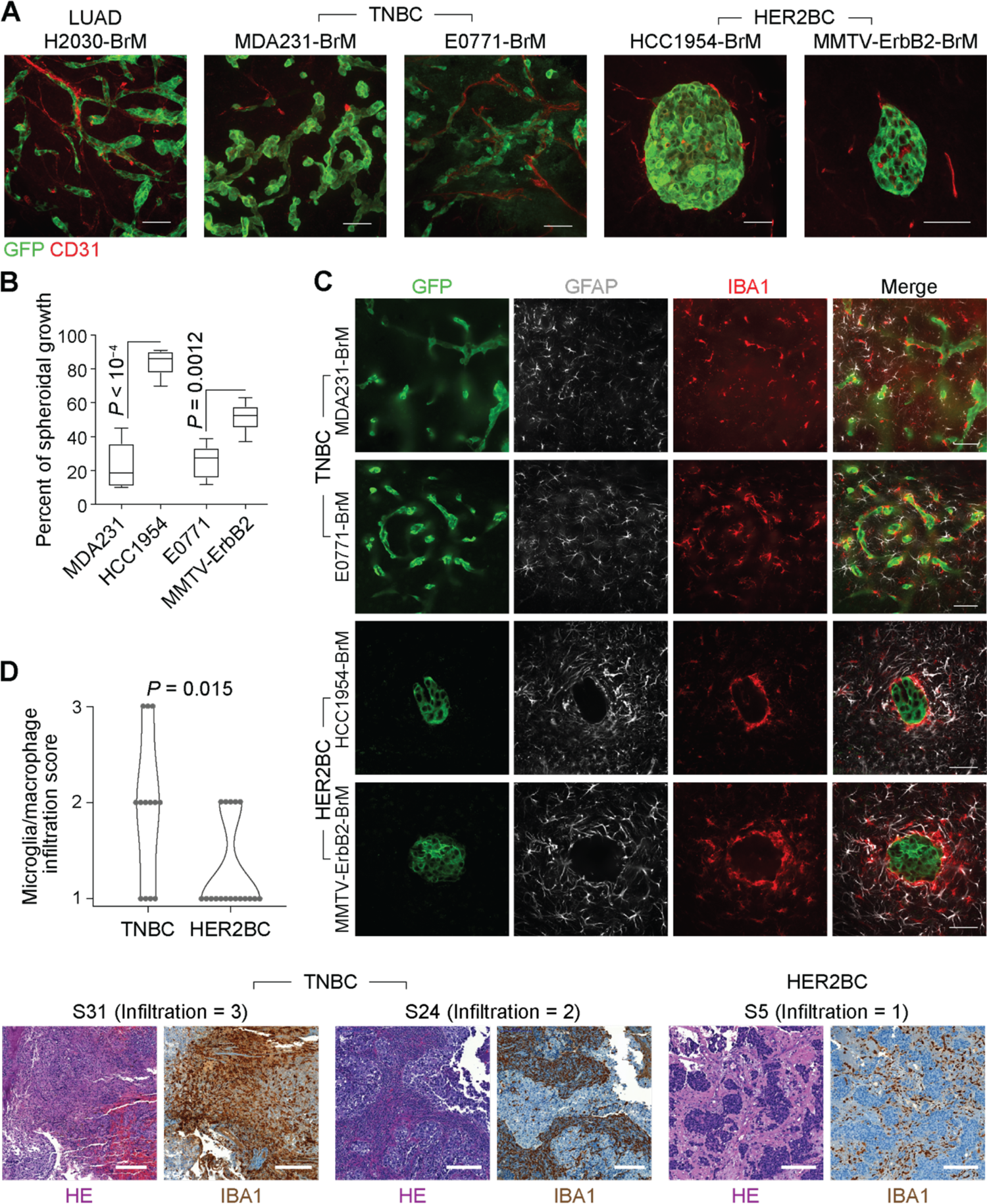
Perivascular and spheroidal brain colonization patterns and stromal interfaces. (A) Representative immunofluorescence (IF) staining of vasculature (CD31+) and tumor lesions (GFP+) formed by brain metastatic derivatives (BrM) of the indicated cell lines. BrM cells were selected by inoculating primary cancer cells into the arterial circulation of mice (human cells into athymic mice, and mouse cells into syngeneic immunocompetent mice), and isolating the subpopulations that preferentially metastasize to the brain parenchyma^11, 94^. Scale bars, 50 µm. (B) Percentage of spheroidal colonies in indicated TNBC and HER2BC brain metastasis models (n = 5-6 mice/group). (C) Infiltrative and segregated patterns of the TME in TNBC and HER2BC brain metastases, respectively. Representative IF staining showing the distribution of astrocytes (GFAP+) and microglia and macrophages (IBA1+) in indicated models. Scale bars, 50 µm. (D) (Top panel) Quantification and (bottom panel) representative H&E staining and immunohistochemistry (IHC) staining of IBA1 (microglia and macrophage infiltration) and associated infiltration scores in brain metastasis tissue samples derived from TNBC patients (n = 13) and HER2BC patients (n = 18). Scale bars, 200 µm.

In contrast to the frequently observed perivascular brain colonization pattern, metastatic HER2BC cells that infiltrate the brain predominantly grow in a spheroidal pattern, as shown with human HCC1954-BrM and mouse MMTV-ErbB2-BrM cells (Figures 1A, 1B, and S1A-S1C). Although not specifically reported by the authors, a spheroidal growth pattern was also apparent in brain metastases from the human HER2BC cell lines JIMT-1 and SUM190^41, 42^. In HCC1954-BrM cells, the spheroidal growth pattern emerged in small clusters consisting of as few as 4 cells (Figure S1C) and remained manifest as clusters grew larger (Figures S1A-S1C). Despite the distinctive spheroidal growth pattern of incipient and established HER2BC colonies, knockdown of *L1CAM* expression using independent short-hairpin RNAs (shRNAs) (Figure S1D) inhibited metastatic growth (Figure S1E), in line with previous finding^13^ of L1CAM being functionally important for a transient stage of vascular cooptive survival after extravasation in HER2BC brain metastasis. Taken together, these results suggested that the metastatic colonization of brain parenchyma followed distinct characteristic patterns, including a diffuse, perivascular pattern in LUAD and TNBC cells previously shown to be mediated by L1CAM, and a previously unreported tight, spheroidal pattern of unknown molecular basis in HER2BC cells.

### Infiltrative and segregated TME in perivascular and spheroidal colony patterns

We next investigated how the distinct TNBC and HER2BC brain colonization patterns spatially interact with astrocytes, microglia, and macrophages, major components of the brain TME^43^. In the vascular-cooptive brain metastatic colonies formed by MDA231-BrM and E0771-BrM TNBC cells, cancer cells were exposed to and co-mingled with astrocytes (identified by GFAP immunofluorescence, IF), and microglia and macrophages (identified by IBA1, IF) (Figure 1C), in agreement with previous reports^15^. In contrast to such infiltrative interface, both astrocytes and microglia/macrophages were largely segregated from HCC1954-BrM and MMTV-ErbB2-BrM spheroidal colonies, where the spatial contact with cancer cells was limited to the periphery of the colonies (Figure 1C). Astrocytes accumulated around the colonies without infiltrating the cancer cell mass. A dense layer of microglia/macrophages enwrapped the colonies, whereas microglia farther away from this layer showed less dense aggregation (Figure 1C).

Similar to these early-stage mouse tumors, in surgically resected brain metastasis tissues from patients harboring large, symptomatic brain metastases, immunohistochemistry (IHC) staining showed a high degree of intermingling of cancer cells with IBA1+ microglia and macrophages in the TNBC cases (n = 13) (Figure 1D and Table 1). Different from the sparsely distributed individual micrometastatic lesions in the mouse model (Figures 1A, 1C, and S1A-S1C), the large lesions of HER2BC cases (n = 18) in patients manifested as aggregated carcinoma cell clusters. However, the absence of infiltrated IBA1+ microglia and macrophages into these clusters was readily apparent (Figure 1D and Table 1), in line with the segregation of microglia and macrophages observed in HER2BC models but not TNBC ones (Figure 1C).

**Table 1.**
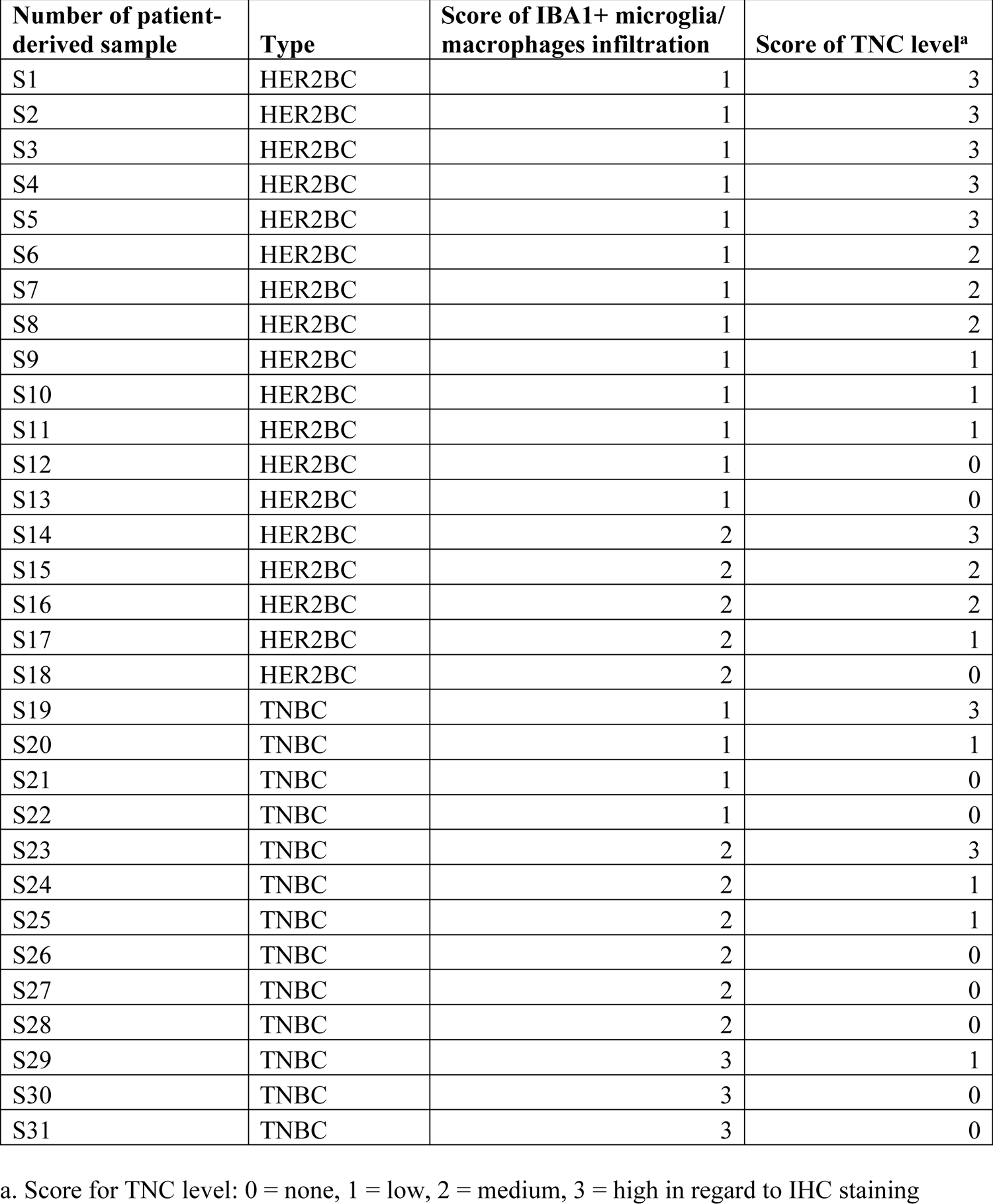
Scoring of IHC-stained patient-derived samples.

### High microglia reactivity in both brain colonization patterns

To characterize the TME of these brain metastatic colonies, we adopted a metastatic niche labeling system^44^ (Figure 2A) that spatially enriches for parenchymal cells situated near the lesions. We engineered MDA231-BrM and HCC1954-BrM cells to constitutively express, in addition to the cell-autonomous GFP, a cell membrane-permeable mCherry protein (sLP-mCherry) secreted outside of the cancer cells and taken up by adjacent cells, acting as a proximity TME label (Figures 2A and 2B). To promote the efficacy of TME labeling, we replaced the mPGK promoter in the original construct^44^ with a stronger *eukaryotic elongation factor 1 alpha 1* (*EEF1A1*) promoter to drive a high level of sLP-mCherry expression. The niche labeling system allowed us to dissociate an entire mouse brain harboring tens to hundreds of MDA231-BrM or HCC1954-BrM early-stage micrometastases, and to use fluorescence-activated cell sorting (FACS) to isolate the GFP+ mCherry+ cancer cells, GFP− mCherry+ TME cells, and unlabeled brain cells, from all lesions pooled together without needing to physically locate each individual lesion prior to dissociation. We harvested and processed the brain tissue samples of the two models in parallel and profiled these samples using single-cell RNA sequencing (scRNA-seq) (Figure 2C). We performed two independent scRNA-seq experiments, and in the second one, included transcription and translation inhibitors during the brain tissue harvesting and homogenizing steps, which may mitigate *ex vivo* perturbation to the glial transcriptome^45^.

**Figure 2.**
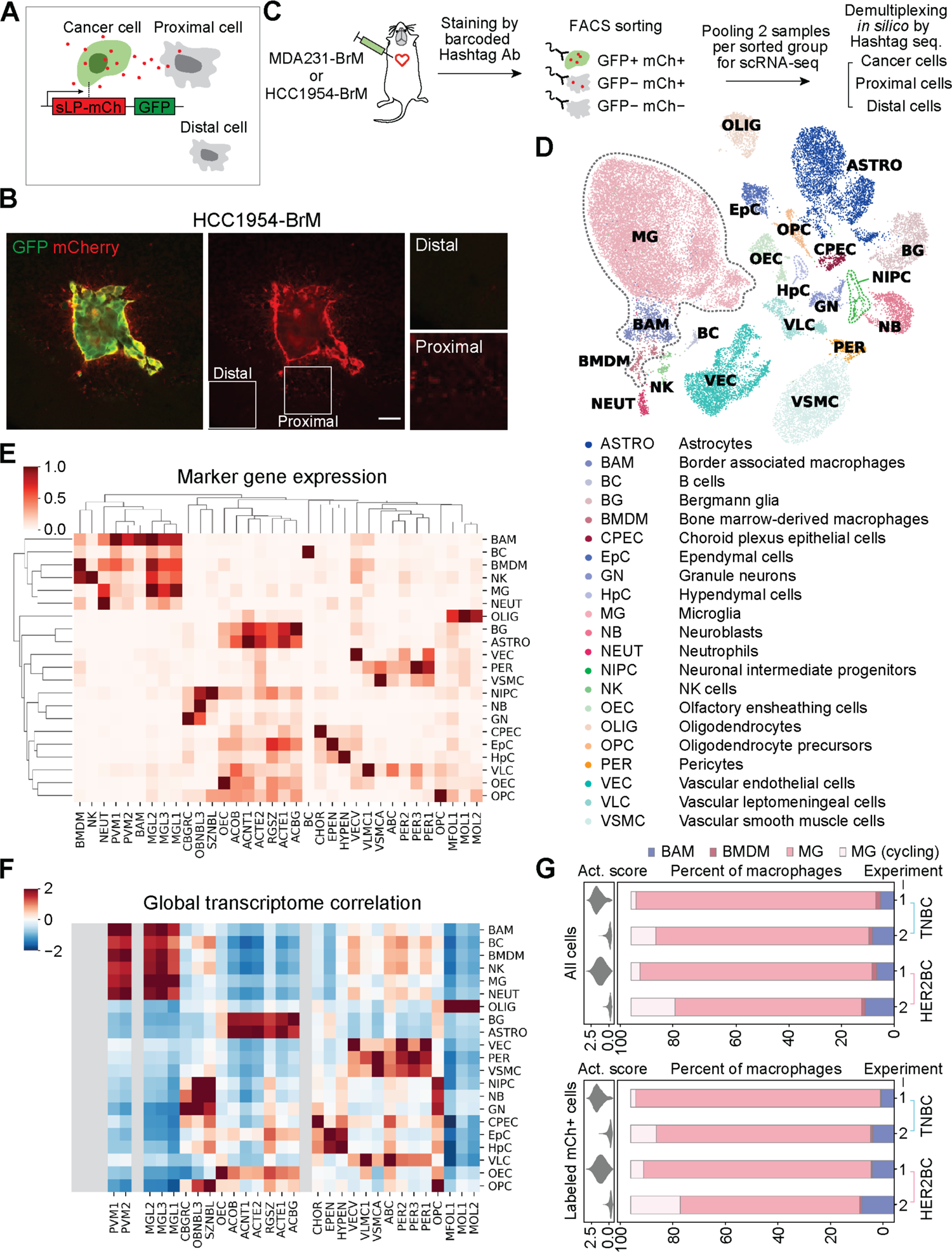
Cellular components of TME by metastatic niche labeling and scRNA-seq profiling. (A) Schematic of the metastatic niche labeling system^44^. Cancer cells co-express GFP and a secreted, lipid-soluble sLP-mCherry protein that penetrates and labels adjacent cells as mCherry+. (B) Representative IF staining showing the labeling HCC1954-BrM HER2BC cells (GFP+, mCherry+), proximal labeled TME cells (mCherry+), and distal unlabeled cells (mCherry−). Scale bar, 50 µm. (C) Schematic of the workflow for profiling single-cell transcriptome of the indicated 3 FACS-sorted groups of cells from dissociated metastases-bearing mouse brains. MDA231-BrM TNBC and HCC1954-BrM HER2BC samples were processed in parallel with cell hashing^95^. Cells from each sample were stained with DNA-barcoded Hashtag antibodies (Ab) recognizing ubiquitous cell surface proteins. Distinct Hashtag sequences allowed pooling cells of two samples for each FACS-sorted group to be sequenced together, and then assigning sequenced cells to their samples of origin based on Hashtag sequences. See STAR Methods for details. (D-F) Calling the cell types of all non-cancer cells from scRNA-seq experiments 1 and 2. (D) UMAP plot of the identified cell populations from 31657 cells. All macrophages, including BAM, BMDM, and MG, are highlighted by dashed gray lines. (E) Clustering showing the population average for the marker genes of reference cell type and state clusters (columns) in each annotated cell population, denoted by the same abbreviated terms as used in (D) (rows, with each row standardized to between 0 and 1). (F) Heatmap of global expression correlation between annotated cell populations (rows) and the cell type and state clusters of reference single-cell transcriptome atlas of the mouse nervous system^46^ (columns, mousebrain.org), z-normalized per row. Rows and columns are organized in identical order with clustermap in (E) to assist visual inspection. Cell types (BMDM, NK, NEUT, BAM, BC) absent in the reference atlas are grayed out. (E, F) Consistent patterns between the marker gene detection (E) and global expression correlation in (F) support robust cell type annotation. See STAR Methods and Table S1 for details of cell type annotation. (G) Tissue dissociation-associated *ex vivo* activation scores (violin plot, left panels) and percentage of the indicated subsets of macrophages (bar plot, right panels) among all macrophages (labeled and unlabeled, top panels) or only labeled macrophages (bottom panels), collected in two independent experiments (1, 2) from the whole brain tissue bearing MDA231-BrM or HCC1954-BrM metastases. MG, microglia computationally identified to be in the G1 phase or not cycling. MG (cycling), microglia inferred to be cycling, given high scores of the S and G2/M phases. BAM, border associated macrophages. BMDM, bone marrow-derived macrophages. Activation scores were computed on the expression of genes that could be further upregulated *ex vivo* by enzymatic dissociation of the brain^45^ (listed in Table S2), and diminished by the addition of transcription and translation inhibitors during brain tissue harvesting and homogenizing in experiment 2.

Using a reference single-cell transcriptome atlas of the mouse nervous system^46^ (mousebrain.org) and additional immune cell type markers complementing this atlas (Figure S2 and Table S1), we identified 21 distinct cell populations among the non-cancer cells, including all labeled and unlabeled cells from both breast cancer models and both experiments (Figures 2D-2F and S2). Given the phenotypic continuums observed in the data, we adopted the Milo framework^47^, which is best suited for identification of subpopulations and cell states that differ in their abundance between conditions, in an unbiased manner without being constrained by predefined cell-type boundaries (clustering). Milo constructed a cell-cell neighbor-graph and performed a statistical comparison of the density between different conditions in neighborhoods across the graph to quantify differential phenotypic shifts between sample conditions (Figure S3A). This analysis revealed that microglia, border associated macrophages, vascular leptomeningeal cells, and oligodendrocyte precursor cells were consistently labeled in both TNBC and HER2BC brain metastases, as demonstrated by the positive log fold-change (logFC) in the abundance of these cells compared to unlabeled counterparts in both scRNA-seq experiments (Figure S3B). The enrichment of these cell populations in the TME (as indicated by positive logFC) suggested that cells of a particular type were present and thus labeled in the TME, and that more importantly, the labeled cells transcriptionally differed from the unlabeled cells of the same type outside of TME if captured, implying that the labeled cells could be reactive to brain metastatic tumor growth despite the different colony architecture and infiltration of the two tumor types. It should be noted that the above transcriptional profiling may fail to efficiently recover certain TME cells (i.e., astrocytes), because such cells poorly survive the tissue dissociation, or they did not take up sufficient sLP-mCherry to be gated as mCherry+ during FACS. Nevertheless, as the metastatic colonies were of a microscopic size (radius from 10 to 1000 µm) and randomly distributed in the brain (Figures S1A and S1B), which render them difficult to dissect or to analyze by currently available spatial transcriptomics technology, the niche labeling system provided a practical, possibly cell type-biased, approach for distinguishing TME cells.

In both MDA231-BrM TNBC and HCC1954-BrM HER2BC models, microglia accounted for more than 90% of all labeled or unlabeled macrophages detected, which also encompassed infiltrated bone marrow-derived macrophages (BMDM) and border associated macrophages (Figures 2G and S3C). This result held in both scRNA-seq experiments, with or without transcription and translation inhibitors during tissue dissociation (Figure 2G). The dominance of microglia also quantitatively agreed with published flow cytometric analysis of tumor-associated macrophages in MDA231 TNBC and 99LN ER+/HER2+ models, which detected ∼10% BMDM and ∼90% microglia^25^. In contrast, the TME composition of surgically resected brain metastasis from patients with breast cancer, lung cancer or melanoma included BMDM constituting more than 50% of tumor associated macrophages^25, 30^. This seeming discrepancy between mouse models and patient samples likely arises from the fact that surgery is performed to resect large, and sometimes pre-treated macrometastatic lesions from advanced stages of metastatic progression that differ from the initial brain colonization stages we focused on in mice. Taken together, the sLP-mCherry system combined with scRNA-seq analysis enabled us to characterize the micrometastatic TME with a spatial resolution that was not feasible with other methods.

### Brain metastases trigger Alzheimer’s disease-associated microglia (DAM) responses

To delineate how microglia react to the brain metastatic colonization, we analyzed the majority non-cycling microglia (i.e., computationally assigned to the G0 or G1 cell cycle phase, Figure S3C, see STAR Methods) of each experiment (Figures 3 and S4) to compare TNBC- and HER2BC-labeled versus unlabeled cells without confounding cell cycle variation. To elucidate potential microglial responses to brain metastatic cells, we first grouped the interconnected phenotypic neighborhoods concordantly enriched in or depleted of both TNBC- and HER2BC-labeled microglia (Figures 3A and 3B) and identified what gene programs were differentially expressed between these two groups of microglia, proximal or distal to the cancer cells, respectively. Interestingly, we found that along with diminishing the expression of basal microglia homeostatic genes (*Hexb*, *Cx3cr1*, *Tmem119*, *Cst3*, *P2ry12*, *blue* in Figure 3C, Tables S2 and S3), brain metastases triggered the Alzheimer’s disease-associated microglial (DAM) responses, as shown by the induction of both global DAM signature genes defined by scRNA-seq transcriptome^48^ (*black*, Figure 3C) and a smaller set of canonical DAM markers as previously highlighted^49^ (*red*, Figure 3C). The DAM phenotype was first identified in Alzheimer’s disease in microglia associated with amyloid-β plaques^48^. The phenotype was later found to be connected, both transcriptionally and functionally, to subsets of microglia from various other developmental and pathological contexts, including in postnatal white matter tracts^50, 51^, amyotrophic lateral sclerosis^52^, and lysolecithin-induced injury^50^, as well as to certain activated macrophages outside of the brain as well, such as the lipid-associated macrophages in adipose tissue^53^. In these various contexts, the core DAM program represents a universal sensor of homeostasis disturbances in microglia, and is accompanied by additional gene responses depending on the specific tissue alteration^49^.

**Figure 3.**
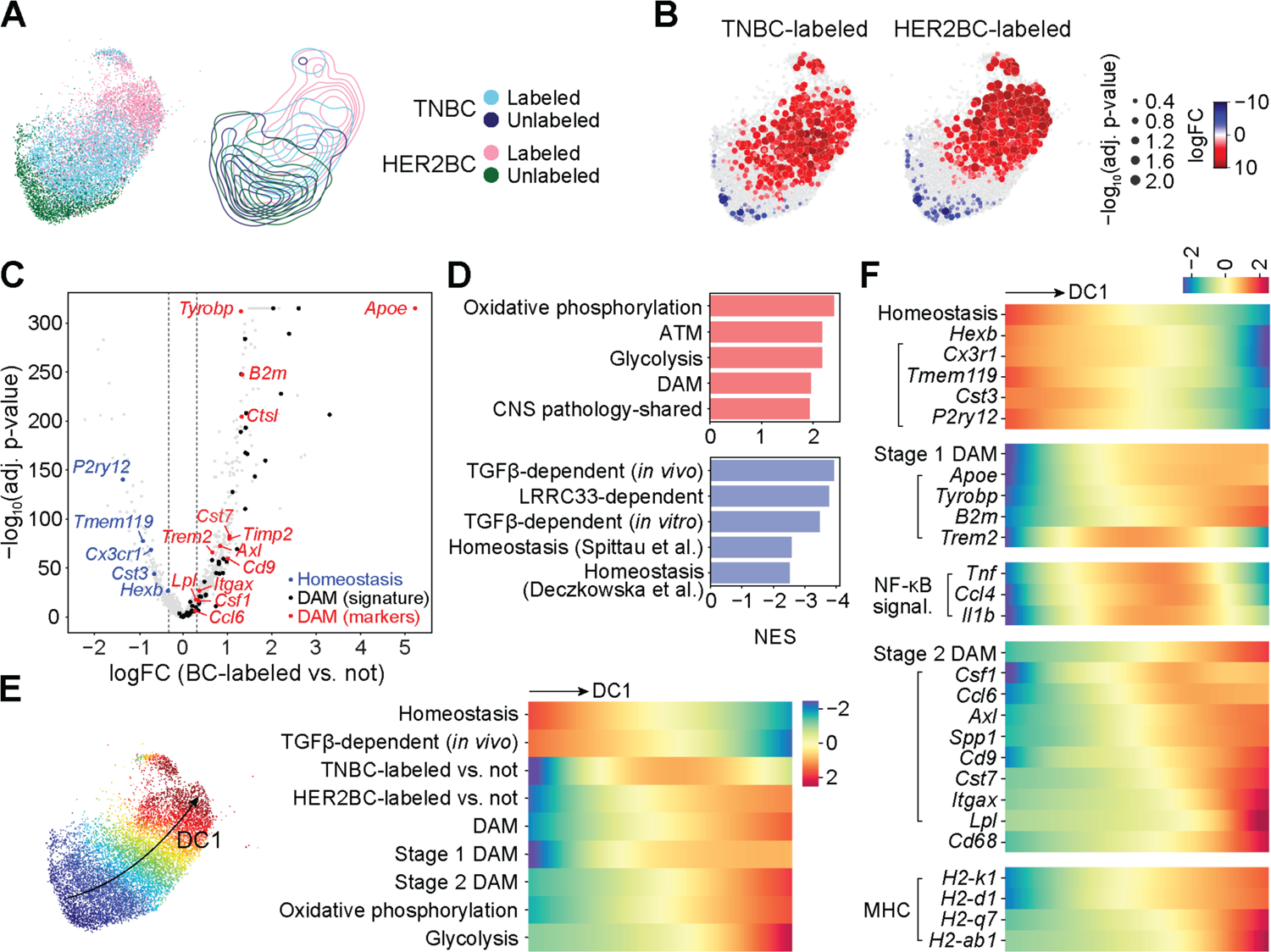
Tumor-associated microglia activate canonical DAM programs. All results computed on the non-cycling microglia (MG) from experiment 1. See Figure S4 for replication of homeostasis-to-DAM transition trends in experiment 2. (A) (Left panel) UMAP embedding of the 4 indicated sources of altogether 11407 cells, and (right panel) contour plots of each source in the embedding. (B) Differential abundance of TNBC-labeled (left panel) and HER2BC-labeled (right panel) cells in reference to unlabeled cells in all phenotypic neighborhoods, overlayed on the UMAP embedding of the index cells of phenotypic neighborhoods. Color and dot size represent the log fold change (logFC) and Benjamini-Hochberg (BH)-adjusted P values of differential abundance, respectively. UMAP embedding of all cells are shown in the background to facilitate visually locating the index cells. (C) Volcano plot of the logFC in gene expression against corresponding BH-adjusted P values, comparing between groups of phenotypic neighborhoods that are concordantly enriched in or depleted of breast cancer (BC) cell-labeled non-cycling microglia, which identified 838 differentially expressed genes (DEGs). logFC thresholds used for calling DEGs are indicated by dashed lines. Sources and lists of genes for DAM^48^ (signature) (*black*), and DAM (markers) (*red*) and homeostasis^49^ (*blue*) are provided in Table S2. (D) Normalized enrichment scores (NES) of the top 5 positively or negatively enriched gene sets among 838 differentially expressed genes (DEGs, Table S3) of labeled non-cycling microglia. See STAR Methods and Tables S2, S4, and S5 for gene set annotations and full GSEA results. (E) (Left panel) UMAP plot showing the first diffusion component (DC1) values computed on all cells by color map; and (right panel) heatmap showing fitted trends of indicated neighborhood-level signature scores and labeled cell enrichment (quantified by the local logFC in abundance in reference to unlabeled cells, logFC, shown in (B)) along the DC1 values of neighborhood index cells (shown in color, left panel) (rows, z-normalized per row across neighborhoods). Homeostasis, stage 1 DAM, and stage 2 DAM marker genes from Ref.^49^; and signatures of TGF-β-dependent expression (*in vivo*), DAM, oxidative phosphorylation, and glycolysis obtained as in (D) (see STAR Methods and Table S2 for details). (F) Similar to (E) heatmap showing fitted trends of neighborhood-level signature scores and gene expression along the DC1 values of neighborhood index cells (rows, z-normalized per row across neighborhoods). Marker genes used to compute the indicated signature scores on their top are indicated by left parentheses.

**Figure 4.**
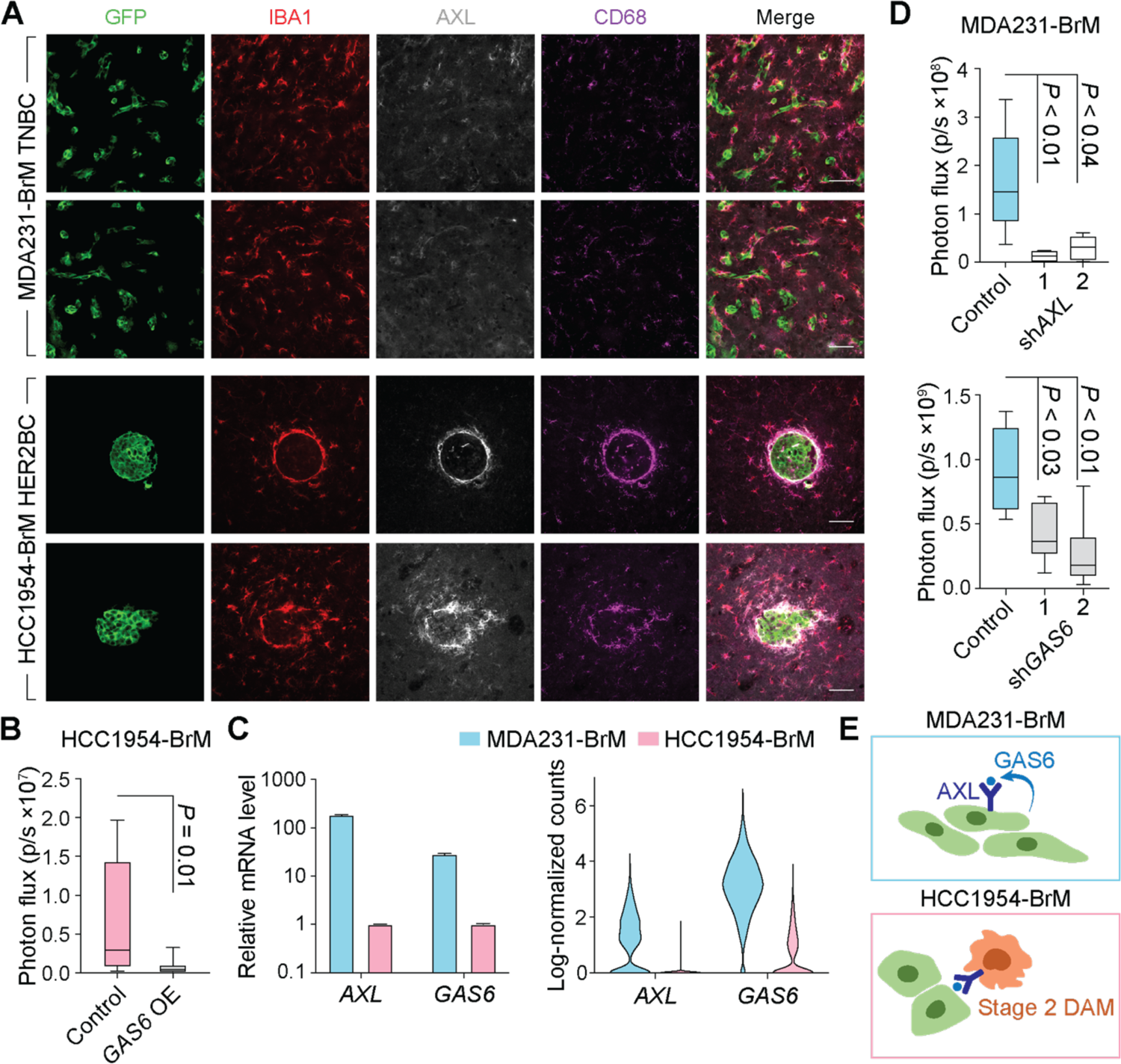
Diverse roles of GAS6/AXL signaling in MDA231 TNBC and HCC1954 HER2BC brain metastases. (A) Representative IF staining of AXL and phagocytosis marker CD68 in IBA1+ microglia/macrophages associated with brain metastatic colonies formed by MDA231-BrM TNBC and HCC1954-BrM HER2BC cells, respectively. Scale bars, 50 µm. (B) Effect of *GAS6* overexpression (OE) in HCC1954-BrM cells on brain colonization measured by *ex vivo* bioluminescence imaging (BLI) of the brain (n = 9 mice/group, 4 weeks post-intracardiac inoculation). (C) *AXL* and *GAS6* expression levels in cancer cells *in vitro* (left panel) and *in vivo* (right panel). (Left panel) Relative mRNA levels of cultured cells measured by qRT-PCR. (Right panel) Log-transformed normalized UMI (unique molecular identifier) counts in single cells, computed on the cancer cells from both experiment 1 and 2 of scRNA-seq analysis. (D) Effect of *AXL* (top panel) and *GAS6* (bottom panel) shRNA knockdown (KD) in MDA231-BrM cells (2 shRNAs per target gene) on brain colonization measured by *ex vivo* BLI of the brain (top panels, n = 6-7 mice/group, 4 weeks post-intracardiac inoculation). (E) Schematic illustrates that GAS6-mediated AXL signaling promotes survival of MDA231-BrM cancer cells (top panel) and mediates phagocytosis of stage 2 DAM in HCC1954-BrM colonies (bottom panel).

We next combined a differential abundance test with diffusion component analysis^54, 55^ to trace shifts in the transcriptional activities of the microglia along a continuous gradient of variation. Gene set enrichment analysis (GSEA) showed that the top diffusion component (DC1), which characterized the major phenotypic variation in the population, strongly correlated with the up- and down-regulation of DAM and homeostasis signatures, respectively (Figure 3D and Tables S3- S5), indicating the homeostasis-to-DAM transition and related gene programs as the primary sources of variation. As consistently observed in both experiments 1 and 2, the homeostasis-to-DAM transition was manifest from the expression trends along DC1: as the expression levels of the DAM signature and the genes comprising it rose, those of homeostasis fell together with a drop in the expression of a TGF-β gene response program – a key regulator of microglia homeostasis^56, 57^ (Figures 3E and S4D). Concomitant with this transition, the relative abundance in TNBC-labeled microglia peaked where the expression of stage 1 DAM marker genes (*Apoe*, *Tyrobp*, *B2m*, *Trem2*) and gene signature started to rise, whereas the relative abundance in HER2BC-labeled microglia showed expression of the phenotypically more advanced stage 2 DAM signature and marker genes (*Csf1*, *Ccl6*, *Axl*, *Spp1*, *Cd9*, *Cst7*, *Itgax*, *Lpl*) (Figures 3F and S4E).

### Differential engagement of two DAM stages

In mouse models of Alzheimer’s disease, microglia transition from homeostasis through an intermediate stage 1 DAM state, marked by induction of stage 1 DAM markers without stage 2 DAM markers, then to stage 2 DAM with additional upregulation of lysosomal, phagocytic, and lipid metabolism genes such as *Axl*, *Cst7*, and *Lpl*^48, 49^. The synchrony in the transitions of identities and transcriptional activities of the microglia along DC1 indicated that in our models, the TNBC metastasis-associated microglia were mostly restricted to the stage 1 DAM state, whereas the HER2BC metastasis-associated microglia largely progressed towards the stage 2 DAM state. This fully developed stage 2 DAM state also displayed enhanced expression of MHC class I (*H2-k1*, *H2-d1*, and *H2-q7*) and class II (*H2-ab1*) genes, and pathways related to oxidative phosphorylation and glycolysis, congruent with the activation of phagocytosis and lipid metabolism in stage 2 DAM^53^ (Figures 3E, 3F, S4D and S4E).

To understand which genes distinguished the stage 1 DAM state from the stage 2 DAM state in brain metastasis, we clustered all genes differentially expressed in labeled microglia by their expression patterns^55^ and identified a cluster (cluster 4 in Table S3) whose expression tracked the differential abundance in TNBC-labeled microglia along DC1. Several genes in this cluster were related to NF-κB-activating inflammatory signals (*Tnf*, *Il1b*, and *Ccl4*) (Figure 3F) observed in aging^50^, brain malignancies^21, 58, 59^, and neurodegenerative disorders^60–62^. *Tnf* and *Il1b* in particular are enriched in the stage 1 DAM of Alzheimer’s disease^63^. As the cytokines TNF, IL-1β, and CCL4 promote tumor growth by directly enhancing cancer cell survival^15^ or by regulating angiogenesis and vascular permeability^64, 65^, their active expression in TNBC metastasis-enriched microglia suggests pro-tumorigenic effects of microglia in this context.

### Disparate roles of GAS6/AXL signaling

IF staining of the receptor tyrosine kinase (RTK) AXL validated the enrichment of stage 2 DAM in HCC1954 HER2BC colonies (Figure 4A). When exposed to certain environmental stimuli, macrophages can increase the level of the receptor tyrosine kinase AXL, and use it to phagocytose apoptotic cells by binding to externalized phosphatidylserine (PtdSer) on these cells via the bispecific AXL and PtdSer ligand GAS6^66, 67^. In Alzheimer’s disease, *AXL* expression is increased specifically in microglia having direct contact with amyloid β plaques, where it mediates the detection and engulfment of amyloid β plaques decorated with GAS6 and PtdSer^68^. Notably, we found that the HCC1954-BrM colonies were demarcated by a rim of AXL+ microglia (Figure 4A). These microglia were phagocytically active as determined by IF staining of the lysosomal marker CD68 (Figure 4A), whose transcript level also tracked that of *AXL* along DC1 (Figures 3F and S4E). Overexpressing *GAS6* in HCC1954-BrM cells (Figure S5A) to increase the concentration of AXL ligand in the TME resulted in a 10-fold reduction in brain metastatic activity (Figure 4B). The enforced surge of GAS6 production potentiated the capacity of surrounding AXL+ stage 2 DAM to eliminate cancer cells in spheroidal HCC1954-BrM colonies. In contrast, the microglia associated with MDA231-BrM colonies were mostly negative for AXL (Figure 4A), in agreement with the scRNA-seq analysis showing limited abundance of stage 2 DAM in TNBC-associated microglia (Figures 3E and S4D).

Aside from being expressed and mediating phagocytosis in macrophages, AXL is also expressed in human TNBC cells^69^ (Figures S5B and S5C), where its activation triggers cancer invasion, survival, and drug resistance through STAT, NF-κB, PI3K, and ERK pathways^70, 71^. Although increased *AXL* expression has been noted in HER2BC cells that undergo EMT^72^, pan-breast cancer cell line analysis of transcriptome (Figure S5B, DepMap database) and proteome (Figure S5C, DepMap database) revealed a specific enrichment of AXL and a significantly higher level of its ligand GAS6 in TNBC cells, compared to HER2BC cells and other types of breast cancer cells. Moreover, *AXL* and *GAS6* transcript levels were positively correlated in TNBC cells (Figures S5B). Previous observational datasets collected through bulk RNA-sequencing (RNA-seq) were not able to distinguish *AXL* or *GAS6* expression from the cancer cell or macrophage populations^73–75^. In agreement with the pan-cell line analysis of parental breast cancer cells, we found that MDA231-BrM cells abundantly expressed *AXL* and *GAS6 in vitro* and *in vivo*. The expression was significantly lower in HCC1954-BrM than in MDA231-BrM, by 200- and 30-fold for *AXL* and *GAS6 in vitro*, respectively (Figure 4C). Knocking down the expression of either gene by independent shRNAs in MDA231-BrM cells inhibited their brain colonization activity by 5- to 10-fold (Figures 4D and S5D), implying that AXL and its ligand GAS6 promoted TNBC brain metastasis by forming an autocrine loop (Figure 4E). In contrast, further reducing the low endogenous expression levels of *AXL* and *GAS6* in HCC1954-BrM cells did not significantly affect brain colonization (Figure S5E and S5F). Overall, the availability of the GAS6 ligand in the TME activated pro-tumorigenic AXL signaling in cancer cells in MDA231-BrM colonies; but in stage 2 DAM in HCC1954-BrM colonies, it activated anti-tumorigenic AXL signaling, yielding differential effects on brain metastases (Figure 4E).

### Heightened expression of extracellular matrix components in brain metastatic HER2BC

To characterize the gene expression patterns that mediate distinct colonization patterns of TNBC and HER2BC brain metastases, we profiled the transcriptome of MDA231-BrM and HCC1954-BrM cells *in situ* using the Flura-seq technique^76^. We engineered cancer cells to express genes encoding cytosine deaminase and uracil phosphoribosyl transferase, which catalyze coupled reactions that convert intraperitoneally administered 5-fluorocytosine, a non-natural pyrimidine, to fluorouridine triphosphate that is incorporated into RNA, and thereby enable metabolic tagging of nascent cancer cell transcripts *in situ* for rapid purification and sequencing^76^ (Figure S6A). Comparing the *in vivo* transcriptome of the two models^77^ revealed high activity of the AKT pathway, a central effector of HER2 signaling, and enriched mesenchymal and extracellular matrix (ECM) assembly signatures in HCC1954-BrM but not MDA231-BrM cells (Figure S6B, see STAR Methods).

To determine whether these *in vivo* signatures relate to brain tropism, we performed RNA expression analysis comparing the parental and BrM derivatives of the HCC1954 and MMTV-ErbB2 HER2BC cells lines, and MDA231 TNBC cells^11^ *in vitro* (Figure 5A). Overrepresentation analysis of the genes differentially expressed in BrM derivatives compared to their corresponding parental cells revealed ECM organization as the top-scoring gene set in HCC1954 and as high-ranking in MMTV-ErbB2, but not in MDA231 (Figure 5B). The *in vitro* transcriptomes of HCC1954 and MMTV-ErbB2 BrM derivatives shared 32 genes that were concordantly upregulated compared to their corresponding parental cells (Table S6). Among them, *COL4A1*, *CST6*, *TNC*, *SEMA7A*, *IL24*, and *S100A7A* encode components of the matrisome, an ensemble of ECM proteins (“core” matrisome, including *COL4A1* and *TNC*), ECM-modifying enzymes, ECM-binding growth factors, and other ECM-associated proteins^78^. Of the two core matrisome members (Figure 5C), tenascin C (*TNC*) is an ECM component of stem cell niches^79^, and collagen type IV alpha 1 (*COL4A1*) is the main collagen type present in basement membranes^80^. The expression levels of both genes displayed a trend of association with relapse in HER2BC patients^81^ (Figure 5D, see STAR Methods for survival analysis of published clinical datasets). Taken together, these results suggested that ECM assembly was correlated with an inherent ability of HER2BC cells to metastasize to the brain.

**Figure 5.**
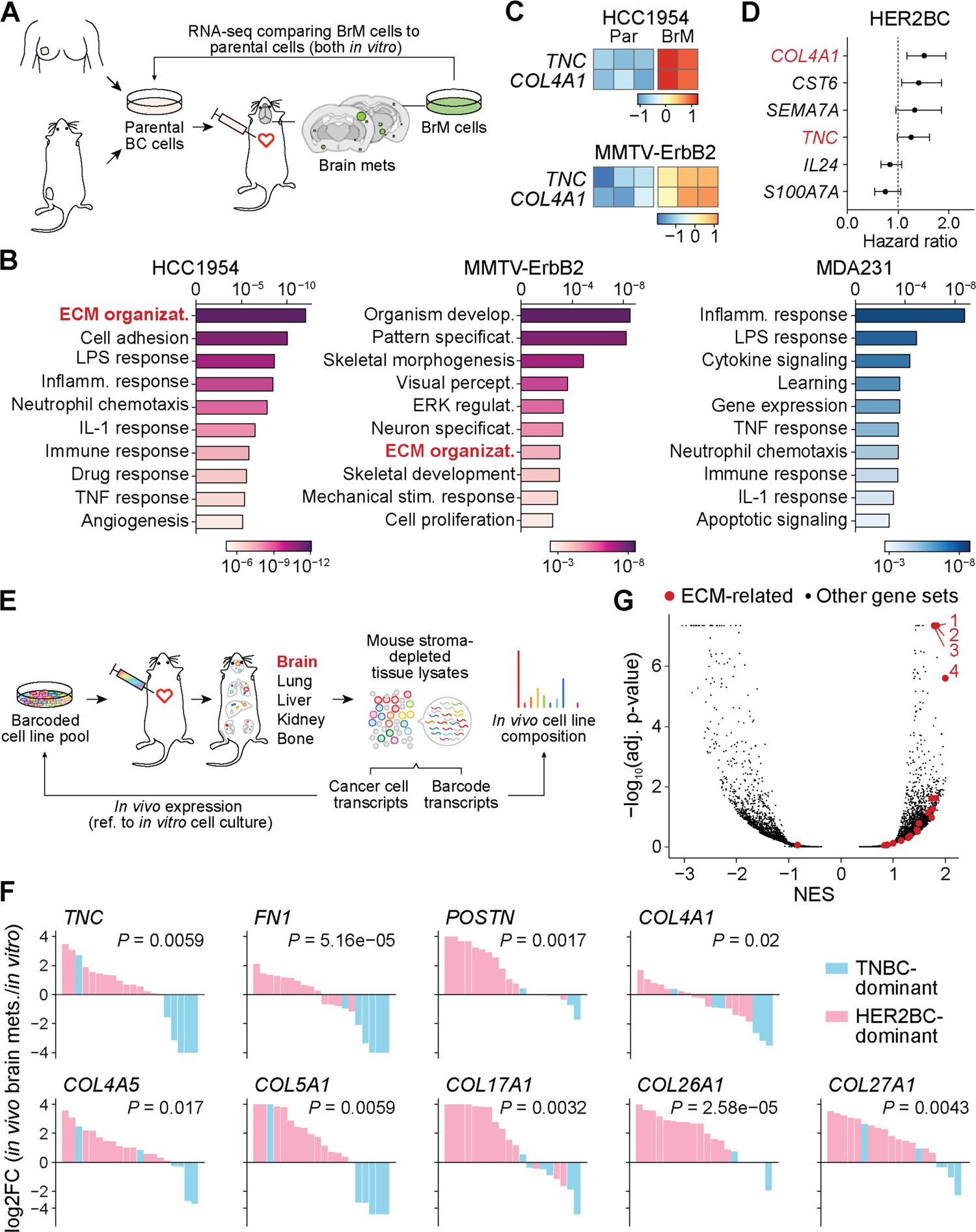
Heightened gene expression of extracellular matrix (ECM) components in HER2BC brain metastasis. (A) Schematic of the workflow of isolating, culturing, and comparing the transcriptome of brain metastatic (BrM) derivatives to the parental breast cancer cell lines they were isolated from. (B) GSEA showing that, in comparison to corresponding parental cell lines, the ECM organization pathway is enriched in the brain metastatic derivatives of HCC1954 and MMTV-ErbB2 HER2BC cells and not in MDA231 TNBC cells. Color shades indicate BH-adjusted P values of normalized enrichment scores. (C) Z-scored expression of the matrisome genes *TNC*, *COL4A1*, *IL24*, and *SEMA7A* that are differentially upregulated in HCC1954 and MMTV-Erbb2 HER2BC brain metastatic derivatives (BrM) in reference to their corresponding parental (Par) cell lines. (D) Forest plots showing the hazard ratio of the expression of matrisome genes *COL4A1*, *CST6*, *SEMA7A*, *TNC*, *IL24*, and *S100A7A* for relapse-free survival of HER2BC breast cancer patients. Gene names in red indicate core matrisome genes. P values from top to bottom are as follows: 0.0012, 0.0135, 0.0894, 0.0641, 0.1649, 0.0889. (E) Schematic of the MetMap workflow using barcoded cancer cell line pools for high-throughput metastatic potential profiling (adapted from Ref.^27^). Relative metastatic potential was quantified by deep sequencing of barcode abundance from tissue. Comparing the transcriptome of *in vivo* brain metastases to that of *in vitro* cell culture per multiplexed cell line pool yielded the log2 fold change (log2FC) of gene expression shown in (F). (F) Relative *in vivo* expression, visualized by the log2FC values shown in a descending order, of the top ECM component genes that are differentially upregulated in the brain metastasis samples composed predominantly of HER2BC cells (pink) than of TNBC cells (blue) (see STAR Methods for statistical analysis of the association between relative *in vivo* expression and percent of HER2BC cells across multiplexed brain metastasis samples). (G) GSEA showing that ECM-related pathways are enriched in multiplexed brain metastasis samples composed predominantly of HER2BC cells (denoted in pink in (F)) than of TNBC cells (in blue in (F)). Top four positively enriched gene sets are Gene Ontology (GO) terms of 1, collagen containing extracellular matrix (GO:0062023), 2, extracellular matrix (GO:0031012), 3, extracellular structure organization (GO:0043062), and 4, extracellular matrix structural constituent (GO:0005201).

To investigate how the molecular features of brain tropism extend to other models of breast cancer, we explored the metastasis map (MetMap) dataset that contains systematically mapped brain metastatic potential and expression patterns of 21 breast cancer cell lines from the Cancer Cell Line Encyclopedia (CCLE, sites.broadinstitute.org/ccle/), including three HER2BC cell lines (HCC1954, HCC1569, JIMT-1) and 18 TNBC cell lines^27^. The cell lines were engineered to express unique 26-nucleotide barcodes and inoculated into NOD-SCID-gamma (NSG) mice as multiplexed pools^27^ (Figure 5E). Most of the brain metastases formed by a pool of cells were found to consist primarily of one subtype of breast cancer as determined by barcode sequencing (Figures S6C and S6D). We could therefore attribute the *in vivo* expression changes from the *in vitro* cell culture, quantified by the log fold change in gene expression (log2FC in Figure 5F), to the dominant cell subtype comprising the brain metastases (Figure S6D), to test whether these changes were associated with TNBC or HER2BC. We found that the expression of multiple core matrisome genes was upregulated or retained *in vivo* in HER2BC-dominant brain metastases, but downregulated in the TNBC-dominant ones (Figure 5F). In addition to *TNC* and *COL4A1*, these genes included those encoding fibronectin (*FN1*) and the tenascin C-binding partner periostin (*POSTN*), both playing vital roles during tissue repair^82^, basement membrane collagens (*COL4A1*, *COL4A5*), transmembrane collagen (*COL17A1*), and ECM collagens (*COL5A1*, *COL26A1*, *COL27A1*). Moreover, GSEA uncovered the ECM-related signatures to be the most highly enriched ones in HER2BC-dominant brain metastases (Figure 5G). Overall, this pan-cancer cell line investigation corroborated that elevated expression of ECM components is closely associated with the brain metastatic potential of HER2BC.

### Cancer cell-derived ECM drives the spheroidal colonization of HER2BC brain metastasis

Using IF staining, we confirmed the presence of TNC, FN1 and POSTN in and around HCC1954 brain metastatic colonies, including the incipient small cell clusters (Figures 6A and 6B). In addition, IHC analysis of patient-derived brain metastasis tissues showed a higher accumulation of TNC in the lesions of HER2BC cases (n = 18) than in those of TNBC cases (n = 13) (Figure 6C).

**Figure 6.**
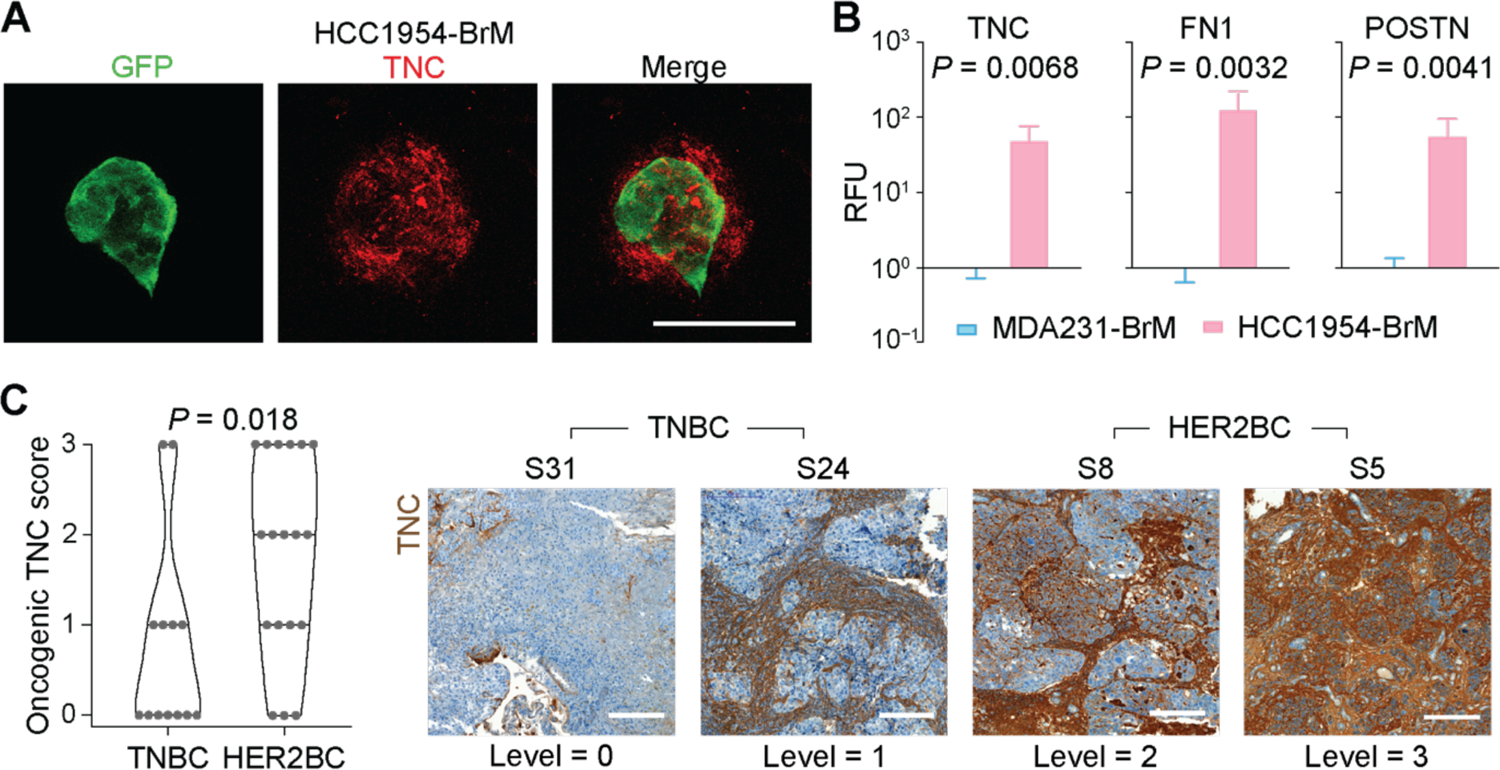
*In situ* validation of tumor ECM deposition in mouse models and clinical samples. (A) Representative IF staining showing the TNC deposition in incipient spheroidal colonies formed by HCC1954-BrM cells 7 days post-intracardiac inoculation. Scale bar, 50 µm. (B) Quantification of the IF staining signal of tenascin C (TNC), fibronectin (FN), and periostin (POSTN) per GFP+ (tumor) unit area in HCC1954-BrM HER2BC and MDA231-BrM TNBC brain metastatic colonies. RFU, relative fluorescence unit. Mean ± SEM. (C) (Left panel) Quantification and (right panel) representative IHC staining and associated TNC level scores (corresponding H&E staining shown in Figure 1D) of TNC in brain metastasis tissue samples derived from TNBC patients (n = 13) and HER2BC patients (n = 18). Scale bars, 200 µm.

To dissect the role of TNC and basement membrane collagens in the formation of HER2BC brain metastatic colonies, we suppressed the expression of *TNC* and *COL4A1* using two different shRNAs in HCC1954-BrM cells and by CRISPR interference (CRISPRi) with two independent sgRNAs in MMTV-ErbB2-BrM cells (Figures S6E and S6F). The suppression of *TNC* or *COL4A1* expression attenuated by two-fold the growth of HCC1954-BrM and MMTV-ErbB2-BrM cells as oncospheres in 3D culture *in vitro* (Figures 7A and 7B), inhibited brain metastasis in both HER2BC models by 5- to 10-fold (Figures 7C and 7D), and reduced the number of incipient HCC1954-BrM colonies formed 21 days post-inoculation by 3-fold (*COL4A1*, *TNC*) (Figure 7E), thereby demonstrating pro-metastatic roles of TNC and COL4A1 in these cells *in vivo*. Although HCC1954-BrM cells (but not MMTV-ErbB2-BrM cells) formed colonies in the lungs when inoculated via the tail vein, the knockdown of *TNC* or *COL4A1* expression did not decrease their lung colonization activity (Figure S6G), suggesting that the pro-metastatic function of TNC and COL4A1 in HER2BC cells was most critical in the brain. Of note, we previously reported that TNC is important for lung metastasis but not brain metastasis of MDA231 cells^83^, indicating that TNC may promote different organotropic metastases depending on the subtype of breast cancer.

**Figure 7.**
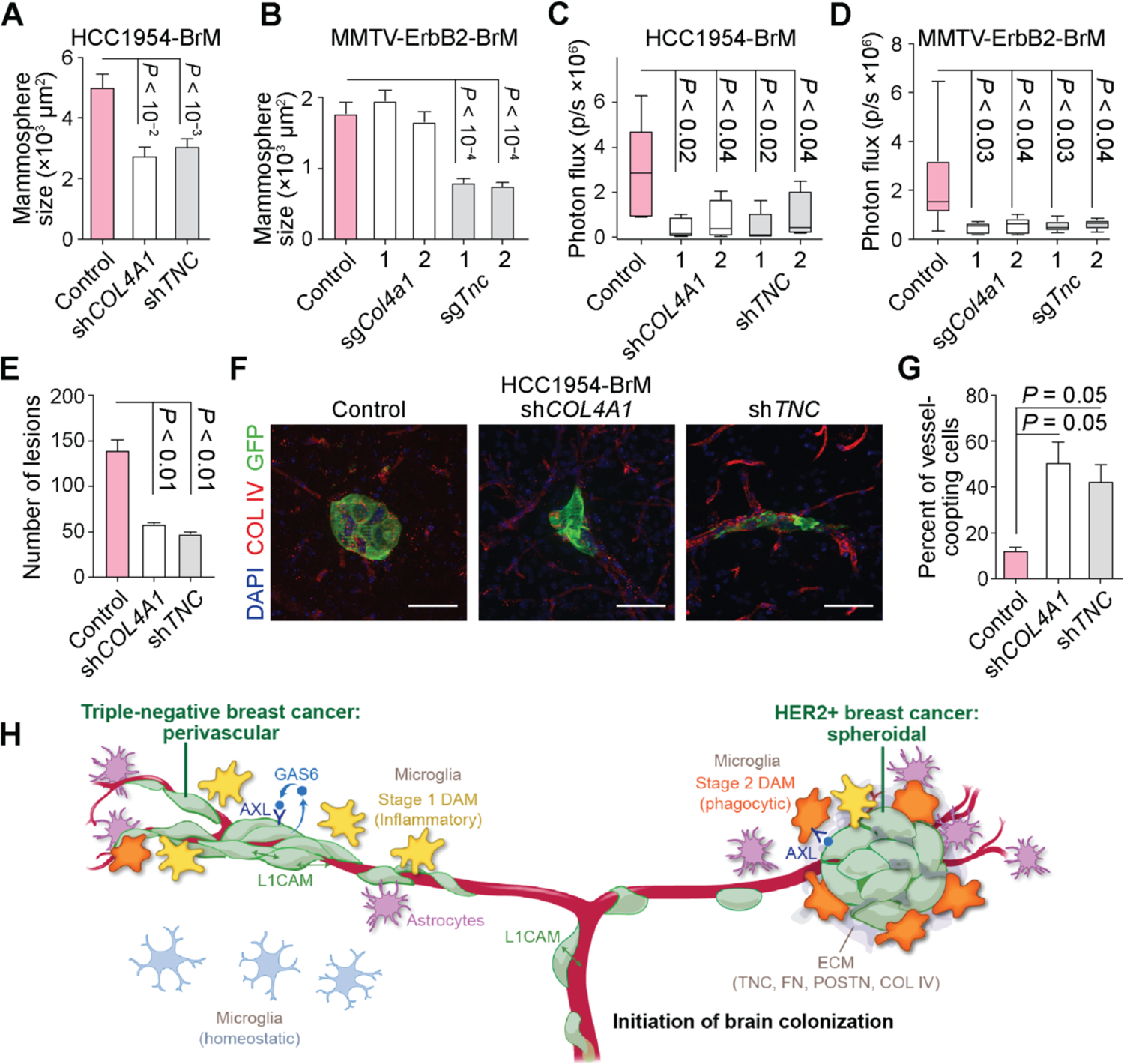
Impact of cancer cell-derived ECM on spheroidal HER2BC brain colonization. **(A-**B) Effect of the suppression of *COL4A1* and *TNC* expression on oncosphere formation (A) by shRNA in HCC1954-BrM cells and (B) by CRISPRi (2 sgRNAS per target gene) in MMTV-Erbb2-BrM cells, measured by the size of colonies after 5 days of growth *in vitro*. n = 100-175 colonies/group. Mean ± SEM. (C-D) Effect of the suppression of *COL4A1* and *TNC* expression on brain colonization (C) by shRNA in HCC1954-BrM cells (n = 5 mice/group, 4 weeks after intracardiac inoculation into athymic mice, 2 shRNAs per target gene) and (D) by CRISPRi in MMTV-Erbb2-BrM cells (n = 5-9 mice/group, 3 weeks post-intracardiac inoculation into FVB/NJ mice, 2 sgRNAs per target gene), both quantified by whole-body BLI. (E-G) (E, G) Quantification and (F) representative IF staining of brain metastatic colonies formed by HCC1954-BrM cells expressing either a control vector or shRNAs that deplete *COL4A1* or *TNC* expression. n = 3 mice/group. Mean ± SEM. Unpaired t test. (G) The percent of vascular coopting cells and (E) the number of colonies were determined 1 and 3 weeks post-intracardiac inoculation, respectively. (H) Schematic illustrates distinctive modes of colonization and stromal interface, various cancer cell-intrinsic mediators of colonization, and the induction of distinct DAM responses during the initiation of brain colonization in TNBC and HER2BC. Copyright © 2023 Memorial Sloan Kettering.

To probe the impact of TNC and COL4A1 on the spheroidal brain colonization architecture of HER2BC, we examined the morphology of single cells and incipient small cell clusters in the HCC1954-BrM model 7 days post-inoculation (Figures 7F and 7G). We observed a preponderance of vascular cooption in the cells with suppressed *TNC* or *COL4A1* expression (Figures 7F and 7G) that preceded the significant reduction in the total number of colonies detected 21 days post-inoculation (Figure 7E). Collectively, these data support a role for TNC and COL4A1 as key drivers of brain-tropic metastasis and spheroidal colony formation in HER2BC.

## Discussion

Here we illuminated novel determinants and tumor-stroma interplay in infiltrating brain metastases by comparing the critical early stages of brain colonization in triple-negative breast cancer (TNBC) and HER2+ breast cancer (HER2BC). These two prevalent subtypes of breast cancer frequently relapse in the brain^4, 5^. However, as we show here, they initiate and establish the colonization with distinct tumor architectures, autocrine growth regulatory mechanisms, and modes of engagement of a highly reactive stroma (Figure 7H). Advances in magnetic resonance imaging for detecting small lesions open the potential for early treatment of brain metastases ^84, 85^; our findings highlight that the TME is an important factor when developing therapeutic strategies to treat metastatic cancers in general.

### Different tumor architectures in the early stages of metastatic brain colonization

The pattern of vascular cooptive growth observed in TNBC was also prominent in lung adenocarcinoma, melanoma, and renal cell carcinoma models^12, 13, 37, 38^, and considered the representative form of brain metastases across multiple primary tumors. The perivascular niche allows cancer cells better access to oxygen and nutrients, and provides anchorage to the vascular basement membrane for survival and outgrowth^86^. Less frequently noted in the brain is the tight, spheroidal growth that HER2BC cells assume soon after infiltration, which is the archetypal growth pattern in a majority of primary tumors^36, 87^. These brain metastatic tumor architectures are respectively associated with infiltrative (TNBC) or segregated (HER2BC) interfaces with the TME. Although phenotypically different, the stromal interfaces of these two major subtypes of breast cancer can both promote brain metastases by facilitating infiltrating cancer cells to draw benefits or evade attack from their TME, as manifested by the impact of GAS6/AXL signaling. The infiltrative and segregated colony phenotypes are predominant in our models of TNBC and HER2BC brain metastasis, respectively, and it is possible that the full range of human disease includes intermediate forms between the infiltrative and segregated phenotypes.

The ECM can promote the seeding and outgrowth of extracerebral metastases^82, 88^, but little is known about its role in brain metastasis^10^. In searching for cancer cell-intrinsic drivers for brain metastasis of HER2BC, we identified a robust ECM deposition program, comprising multiple interacting collagens (type IV collagen) and glycoproteins (tenascin C, fibronectin, and periostin). Type IV collagen is the major constituent of the basement membrane that provides structural and signaling support to surrounding tissue^80^. Tenascin C, fibronectin, and periostin can directly bind to each other and with integrin cell adhesion receptors^89–91^ to induce stemness-related signaling pathways in development and wound healing^82, 83^. We show that *TNC* or *COL4A1* knockdown shifts incipient lesions into the perivascular mode found in HER2BC cells shortly after extravasation and reduces overall metastatic growth. These concurrent changes imply that prolonged post-extravasation spreading on the vasculature is detrimental to the survival of HER2BC cells, and that the deposition of ECM components enables these cells to exit the transient L1CAM-mediated vascular cooption^13^ and adopt a spheroidal growth mode to colonize the brain. In short, an autocrine ECM program supports the interlinked survival and architecture of HER2BC brain metastasis.

### Overlapping Alzheimer’s disease-like microglia responses in different tumor architectures

Our scRNA-seq analysis revealed non-identical, albeit overlapping, disease-associated microglia (DAM) responses the microglia associated with both TNBC and HER2BC brain metastases. Despite distinct colonization patterns and spatial interfaces with the stroma, MDA231 TNBC and HCC1954 HER2BC brain metastases activate – to various degrees and stages – conserved DAM responses originally defined in Alzheimer’s disease, with the inflammatory stage 1 DAM and phagocytic AXL+ stage 2 DAM enriched in TNBC and HER2BC colonies, respectively. Expression of a limited number of DAM marker genes has been noted in the bulk-averaged transcriptome of all microglia from a mouse model of LUAD brain metastasis^23^. However, the DAM phenotype and its specific stages are a function of broad gene expression programs, which we examine in spatially enriched TME at single-cell resolution to dissect the reaction of metastasis-associated microglia. A pan-cancer study of surgically resected human brain macrometastases was recently reported^32^. Our analysis of this dataset revealed heterogenous enrichment of various DAM genes in tumor-associated macrophages, including genes annotated as canonical markers (like *APOE*, *TREM2*, *SPP1*, and *AXL*), and also those that have been reported by multiple DAM studies^92, 93^ (such as *APOC1*, *C1AQ*, and *IL1B*), all of which are significantly upregulated in the metastasis-associated microglia in our BrM models (Table S3). Such expression patterns identify the DAM phenotype as a common feature of the TME in brain metastasis, conserved between patients and mouse models, and shared across stages and primary cancer types.

Our findings on the GAS6/AXL axis provide an example of the tumor-microglia interplay that carries therapeutic implications. This signaling axis causes varied systemic effects depending on the cell types expressing the AXL receptor and the tumor architectures and corresponding stromal interfaces that influence the availability of its ligands. The close contact of TME cells with perivascular TNBC colonies might facilitate access to stroma-derived GAS6 that supplements cancer cell-derived GAS6 to trigger pro-survival GAS6/AXL signaling in cancer cells. In contrast, the stromal segregation in spheroidal HER2BC colonies may serve as a protective barrier limiting the exposure of cancer cell mass to the canonical AXL+ phagocytic stage 2 DAM bridged to cancer cells by stroma-derived GAS6. Treating brain metastases in these TNBC and HER2BC cases may require different options: inhibiting GAS6/AXL signaling may suppress brain metastases that resemble TNBC cases, whereas enhancing GAS6/AXL signaling and may suppress brain metastases in HER2BC cases.

Future studies will be required to systematically unravel the molecular mechanisms by which different cancer cell types and DAM stages influence each other, to guide the design of therapeutic interventions that function by activating or reinforcing the DAM stage that exerts detrimental effects on tumor growth or by blocking cancer cells from receiving growth benefits from the DAM stage induced. This information will be particularly relevant for developing new treatment strategies for brain micrometastatic disease.

## Acknowledgements

We thank the MSKCC Single-Cell Analytics Innovation Lab, Integrated Genomics Operation, Flow Cytometry Core Facility, Molecular Pathology Core Facility, Huiyong Zhao from Antitumor Assessment Core Facility, and Catherine Bibby for their technical assistance. We thank Terry Helms from Design and Creative Services for generating the Figure 7H illustration. We thank Griffin Hartmann, Thomas Tammela, Vijay Yarlagadda, Mollie Chipman, and Russell Kunes for help with experimental and computational aspects, and Adrienne Boire, Joao Xavier, Jose Reyes, Jing Hu, Samuel Rose, and Russell Kunes for comments on the manuscript. This work was supported by NIH grants P01-CA129243 (J.M.), U54-CA209975 (J.M., D.P.), and R01-DK127821 (A-K.H.), P30-CA008748 (Memorial Sloan Kettering Cancer Center), Damon Runyon Quantitative Biology Fellowship (S.G.), the Center for Experimental Immuno-Oncology Fellowship (S.H.S), the Alan and Sandra Gerry Metastasis Research Initiative (D.P., J.M.), and Cycle for Survival (J.T.M., A-K.H.). D.P. is an HHMI Investigator.

## Author Contributions

S.G., D.G.M., L.T., D.P., and J.M. conceived studies, designed experiments, interpreted results, and wrote the manuscript; S.G., D.G.M., S.H.S., L.T., and H.B. performed experiments and analyzed data; X.J. analyzed the MetMap dataset; S.G. and D.P. analyzed scRNA-seq data; J.T.M. and A-K.H. guided collecting and analyzing the iDISCO data; P.A., E.S., W.C., and N.S. analyzed the DepMap dataset; E.B. and T.A.B. oversaw the tissue processing and histopathological data interpretation of clinical samples.

## KEY RESOURCES TABLE

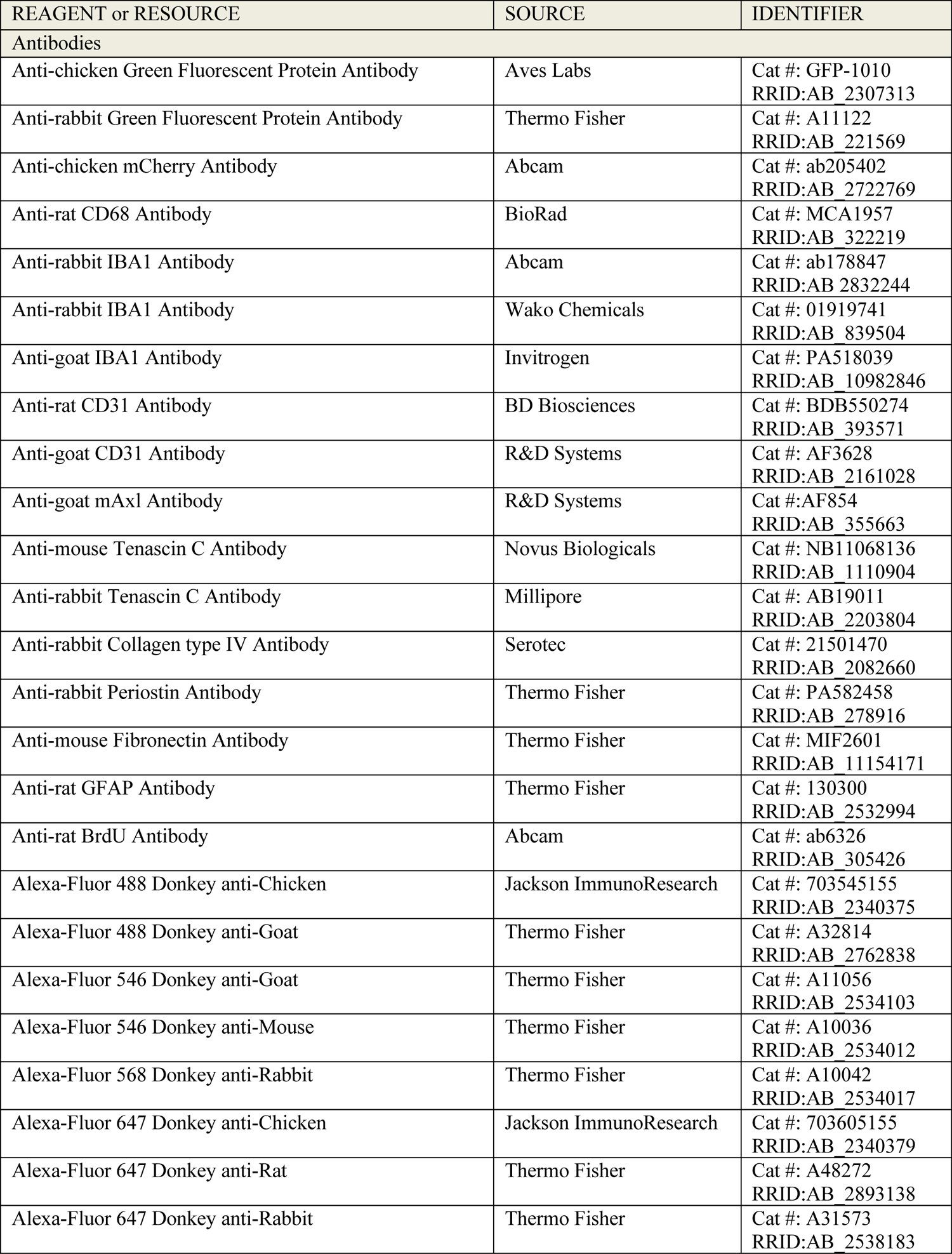
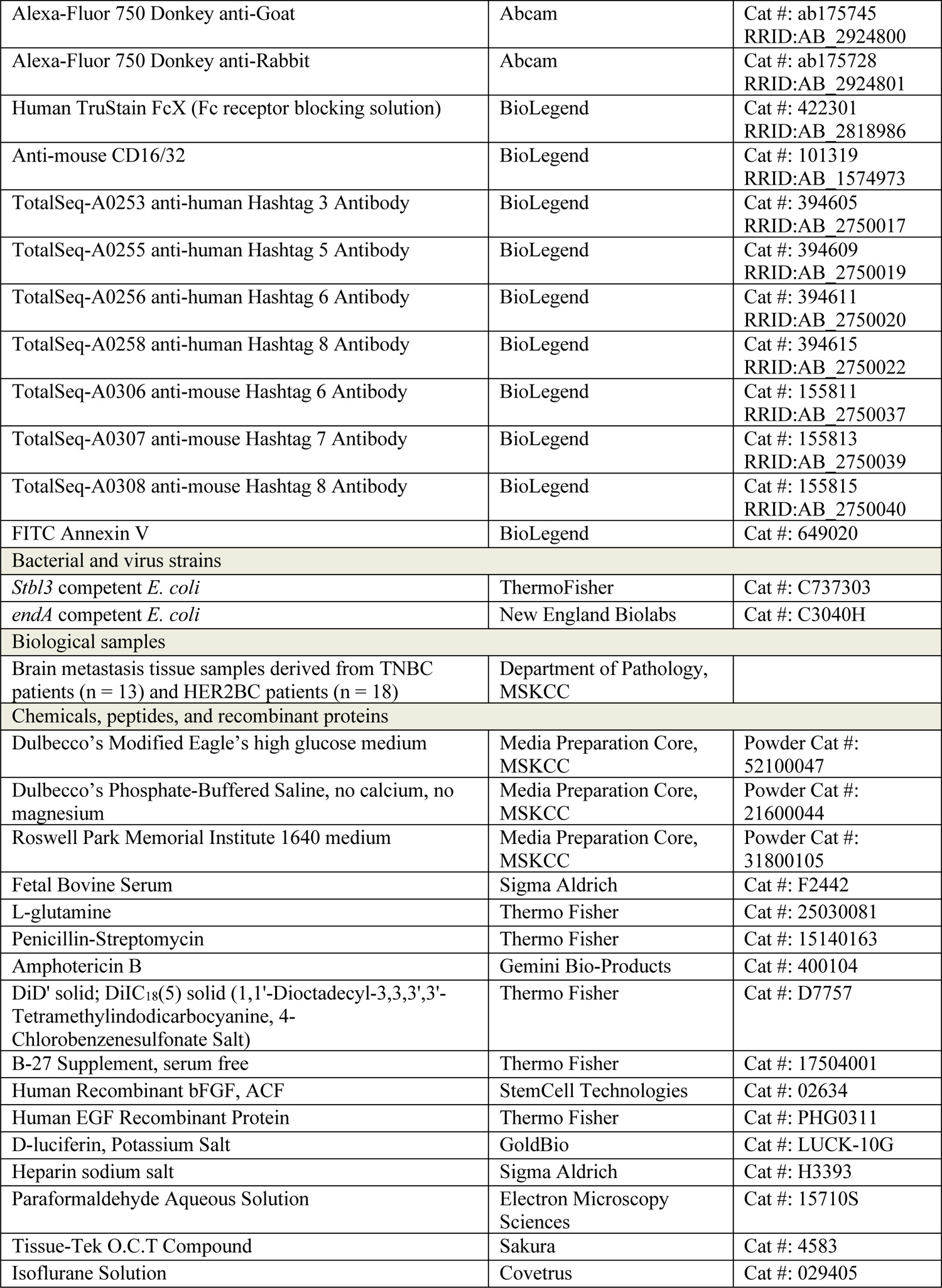
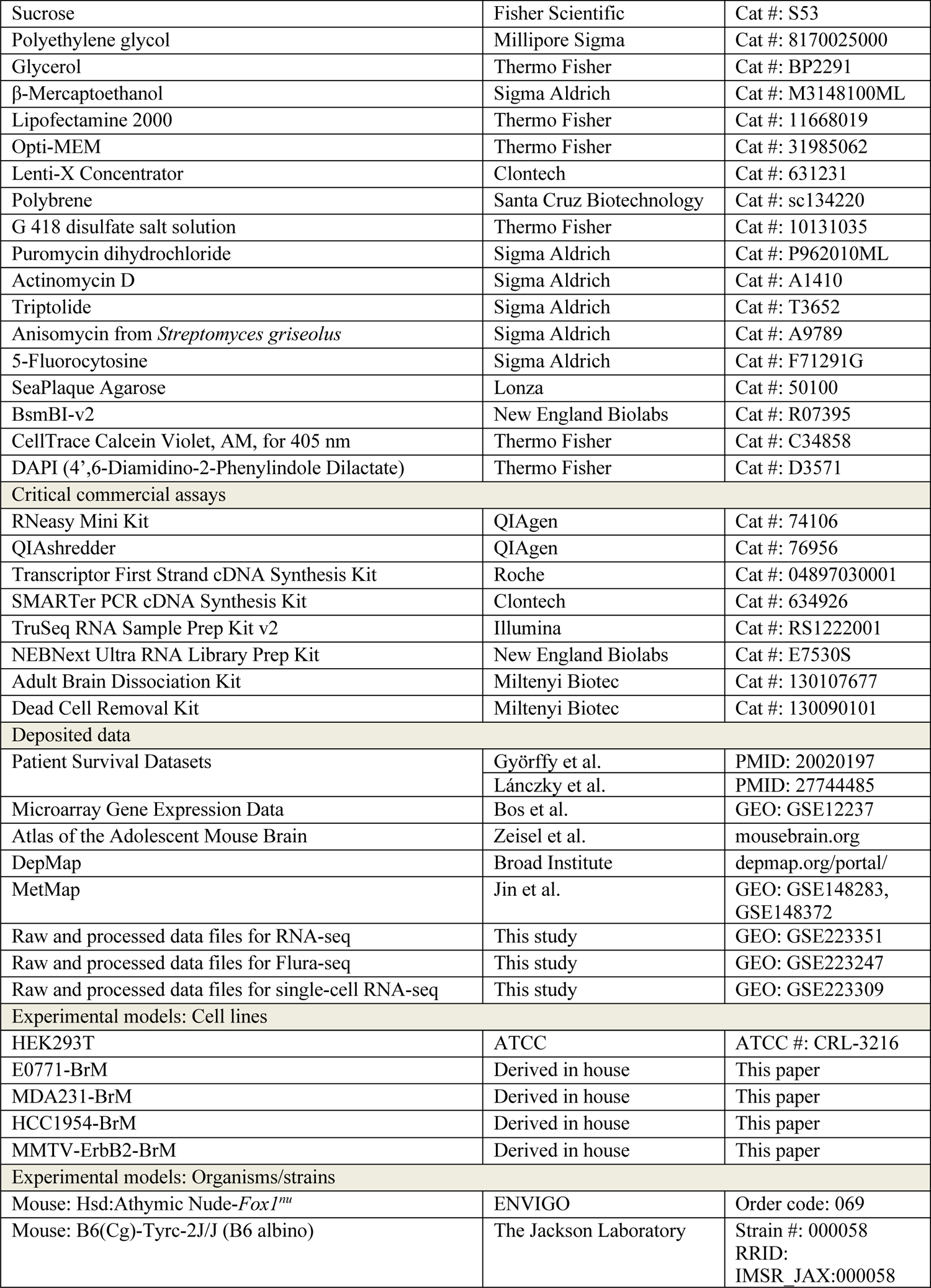
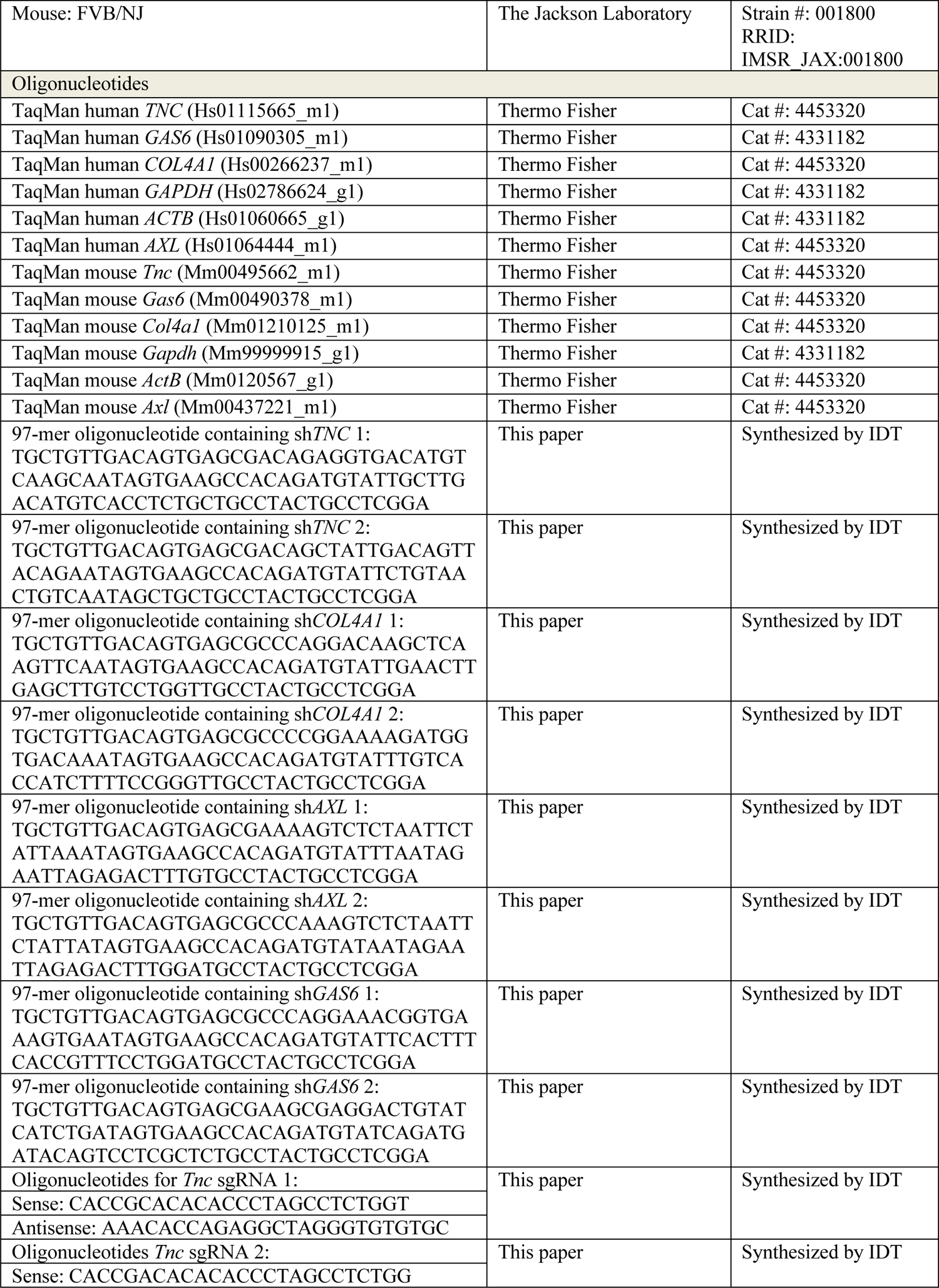
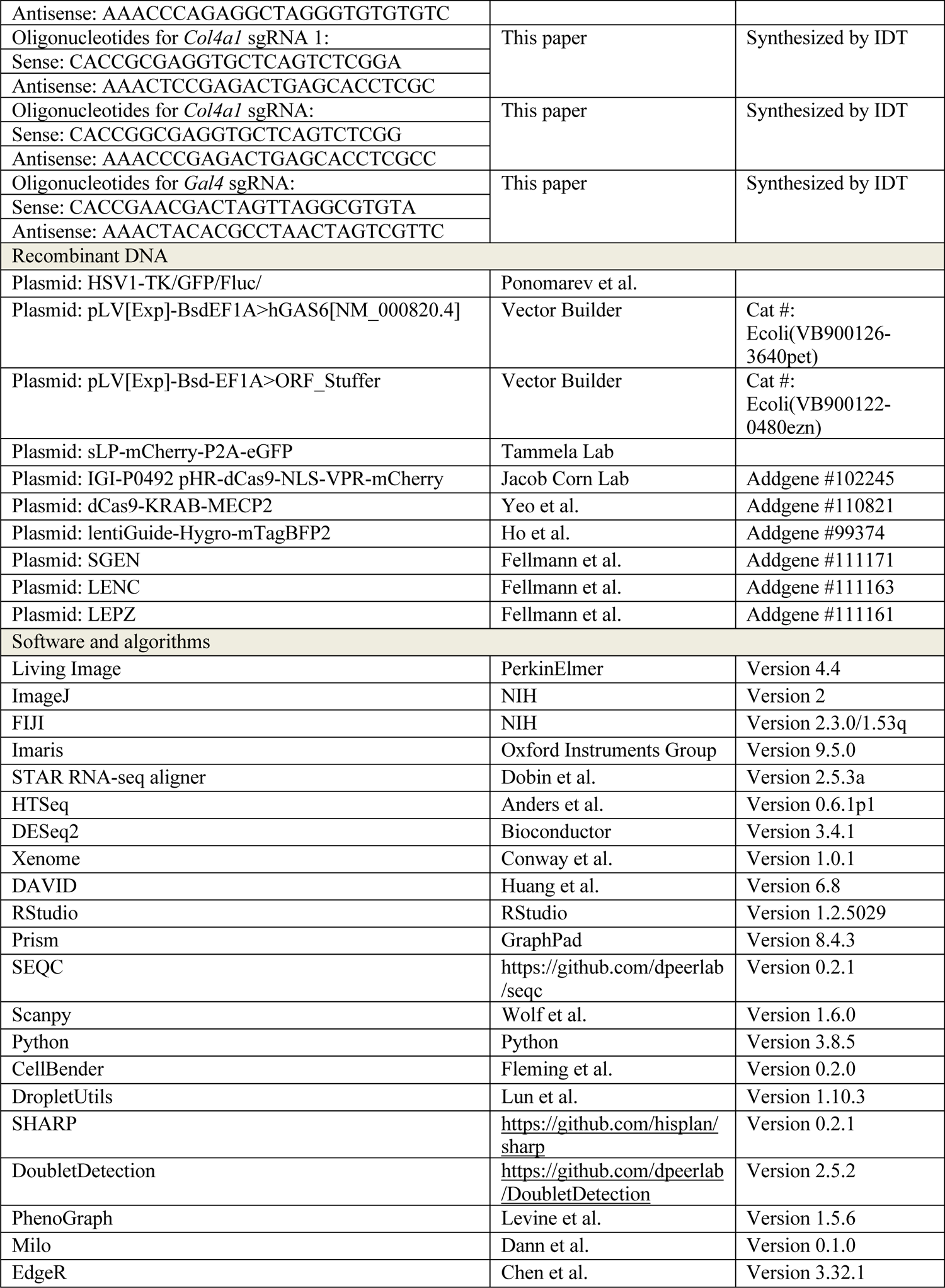
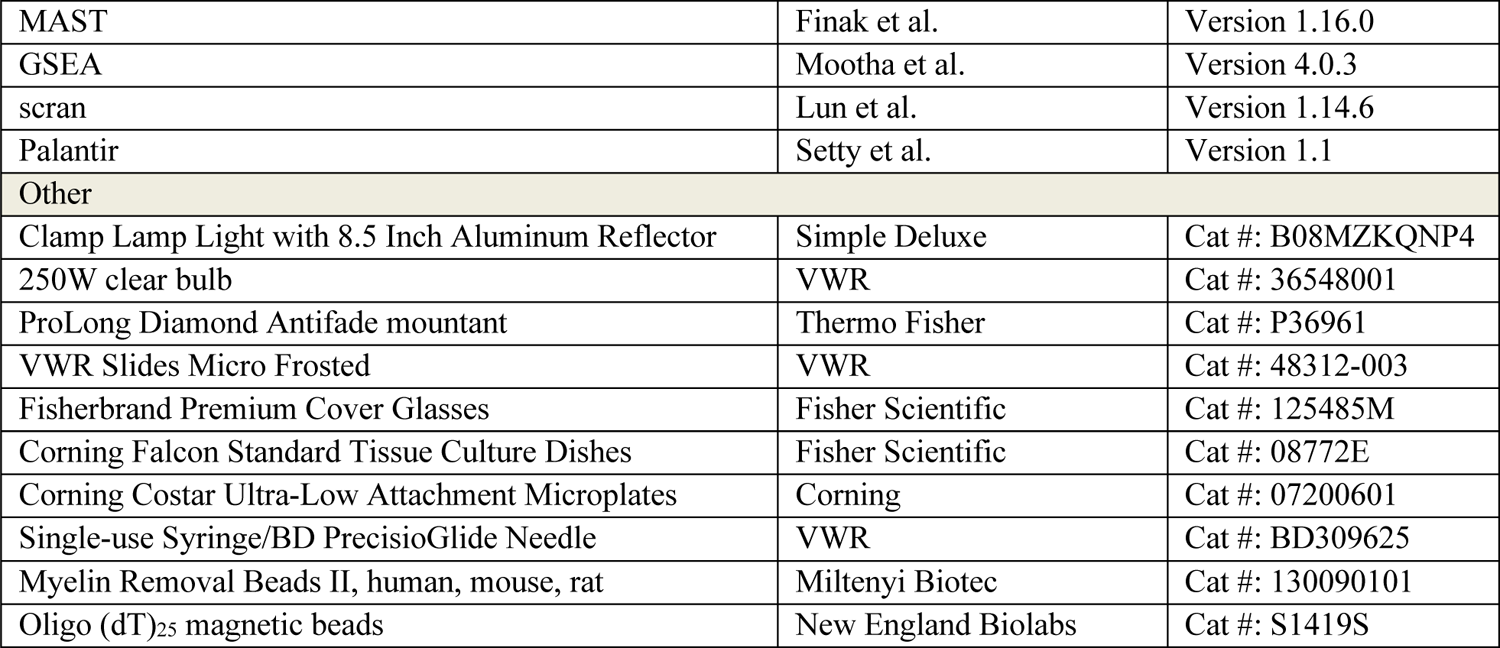

## STAR ★ METHODS

### RESOURCE AVAILABILITY

### Lead contact

Further information and requests for resources and reagents should be directed to and will be fulfilled by the lead contact Joan Massagué.

### Materials availability

All unique reagents generated in this study, including plasmids and cancer cell lines, are available from the lead contact with a completed Materials Transfer Agreement.

### Data and code availability

Bulk and single-cell RNA-seq data have been deposited in the Gene Expression Omnibus database and are publicly available as of the date of publication. Accession numbers are listed in the key resources table. R and Python codes for conducting the scRNA-seq analysis will be uploaded to GitHub (github.com/dpeerlab). All software programs used for analyses are publicly available and listed in the key resources table. Microscopy data and any additional information required to reanalyze data reported in this paper are available from the lead contact upon request.

### EXPERIMENTAL MODEL AND SUBJECT DETAILS

#### Cell lines

MDA231 and MMTV-ErbB2, including parental cells and BrM derivatives, as well as HEK293T cells were cultured in Dulbecco’s Modified Eagle’s (DME) high glucose medium (Media Preparation Core, MSKCC, Cat: 52100047). HCC1954 and E0771, including parental cells and BrM derivatives, were cultured in Roswell Park Memorial Institute (RPMI) 1640 medium (Media Preparation Core, MSKCC, Cat: 31800105). Both media were supplemented with 10% fetal bovine serum (FBS) (Sigma Aldrich, Cat: F2442), 2 mM L-glutamine (Gln) (Thermo Fisher Scientific, Cat: 25030081), 100 IU/mL penicillin/streptomycin (P/S) (Thermo Fisher Scientific, Cat: 15140163), and 1 µg/mL amphotericin B (Gemini Bio-Products, Cat: 400-104). All parental cells were obtained from ATCC and were female in origin. All cells were grown in a humidified incubator at 37 °C with 5% CO_2_ and verified to be mycoplasma-free on a monthly basis.

#### Animals

All animal experiments were conducted in accordance with a protocol approved by the Memorial Sloan Kettering Cancer Center (MSKCC) Institutional Animal Care and Use Committee (IACUC). Athymic nude (Hsd:Athymic Nude-*Foxn1^nu^*, ENVIGO, order code: 069) female mice aged between 5-7 weeks were used for metastatic colonization assays of MDA231-BrM cells and HCC1954-BrM cells. B6(Cg)-Tyrc-2J/J (B6 albino mice, strain #: 000058) and FVB/NJ (strain #: 001800) female mice aged between 5-7 weeks, both from The Jackson Laboratory, were used for metastatic colonization assays of E0771-BrM cells and MMTV-ErbB2-BrM cells, respectively.

#### Clinical samples and immunohistochemistry

All human tissues were obtained under MSKCC Institutional Review Board biospecimen research protocol 15-101. All patients provided pre-procedure informed consent. The archival formalin-fixed, paraffin-embedded (FFPE) brain metastasis of HER2BC (18 cases) and TNBC (13 cases) clinical tissue blocks used for immunostaining were identified by database search and chart review. IHC was performed by the MSKCC Molecular Cytology Core using antibodies against Tenascin C (Millipore, Cat: AB19011, 0.5 µg/mL) and IBA1 (Abcam, Cat: ab178847, 0.2 µg/mL). Tissue processing and histopathological data interpretation were overseen by an expert breast cancer pathologist (E.B.).

## METHOD DETAILS

### Brain metastatic cell isolation

Brain-tropic metastatic derivatives (BrM) of cell lines MDA231, HCC1954, and MMTV-ErbB2 were established as previously described^11, 12, 94^. BrM derivatives of cell line E0771 were generated following the same procedure via two cycles of *in vivo* selection of the parental cells for their preferential metastasis to the brain in background-matched FVB/NJ mice (The Jackson Laboratory, strain #001800). Briefly, E0771 cells were transduced with lentivirus expressing the triple-fusion reporter encoding herpes simplex virus thymidine kinase 1 (HSV1-TK), GFP and firefly luciferase^96^. 1.0 × 10^5^ parental cells were injected into the left ventricle of anesthetized 5-7 week-old FVB/NJ mice (The Jackson Laboratory, strain #: 001800) in a volume of 100 µL. Tumor development was monitored by weekly bioluminescence imaging (BLI) using the Xenogen IVIS-200 system (PerkinElmer). We harvested and dissociated brains with positive BLI signal into single-cell suspension, and isolated GFP+ cells by FACS sorting to be subject to a second round of *in vivo* selection to obtain BrM derivatives.

### Animal studies

Brain metastatic colonization assays were performed as follows: mice were anaesthetized by 100 mg/kg ketamine and 10 mg/kg xylazine. 1.0 × 10^5^ BrM cells resuspended in 100 µL ice-cold phosphate buffered saline, no calcium, no magnesium solution (PBS, Media Preparation Core, MSKCC, Cat: 21600044) supplemented with 2% FBS were injected into the left ventricle of the mice with a 26G × 3/8” needle attached to tuberculin syringe (VWR, Cat: BD309625). During the course of colonization assays, metastatic burden was monitored by non-invasive bioluminescence imaging (BLI) *in vivo* using the Xenogen IVIS-200 system. Specifically, mice were anaesthetized in an induction chamber connected to isoflurane (Covetrus, Cat: 029405), and imaged within 2-4 minutes after retro-orbital injection of 1.5 mg D-luciferin (GoldBio, Cat: LUCK-10G) dissolved in 100 µL PBS. After a designated period of time post-injection indicated in the figure legends for individual experiments, mouse brains were collected for histological analysis. Briefly, mice were imaged by BLI *in vivo* as described above (see **Brain metastatic cell isolation**), euthanized by CO_2_, and transcardially perfused with 10 mL PBS containing 1.5 mg D-luciferin potassium salt and 10 mg/L heparin sodium salt (Sigma Aldrich, Cat: H3393). Whole brains were immediately isolated, imaged by BLI *ex vivo*, and subsequently incubated with rotation at 4 °C first in 4% paraformaldehyde (PFA) (Electron Microscopy Sciences, Cat: 15710-S) for 24 hours and then in 30% (w/v) sucrose (Fisher Scientific, Cat: S53) in PBS for 48 hours, with three PBS washes in between. PFA-fixed, sucrose-preserved brains were embedded into Tissue-Tek O.C.T. compound (Sakura, Cat: 4583), mounted onto the platform of a sliding microtome (Thermo Fisher Scientific, Cat: HM450), frozen to −30° C. 80 µm-thick slices were sectioned, and serially stored in ten 2 mL volume centrifuge tubes (USA Scientific, Cat: 14209704) at −20 °C in anti-freezing solution, containing 30% polyethylene glycol (Millipore Sigma, Cat: 8170025000) and 30% glycerol (Thermo Fisher, Cat: BP2291) in PBS. Lung colonization assays were similarly performed except that BrM cells were injected into the lateral tail vein of the mice to initiate lung colonization. In both brain and lung colonization assays, BLI signal was measured using the ROI tool in Living Image software (PerkinElmer, version 4.4)

### Oncosphere culture

Single-cell suspensions of HCC1954-BrM cells and MMTV-ErbB2-BrM cells were plated in ultra-low attachment plates (Corning, Cat: 07200601) at a density of 1.0 × 10^5^ cells/mL in corresponding culture media supplement with 1X B-27 (Life Technologies, Cat: 17504-001), 20 ng/mL human recombinant bFGF (StemCell, Cat: 02634), 20 ng/mL human EGF recombinant protein (Life Technologies, Cat: PHG0311), and 100 IU/mL P/S. Cells were cultured for 5 days, and imaged with EVOS Cell Imaging Systems (Thermo Fisher). ImageJ (version 2) was used to quantify the diameter of the oncospheres.

### Immunofluorescence (IF) staining and imaging of free-floating tissue sections

Tissue sections archived in anti-freezing solution were washed thoroughly in PBS three times to remove residual cryoprotectant. Sections were permeabilized with two washes of PBS-T, that is, PBS supplemented with 0.25% Triton X-100 (Fisher Scientific, Cat: AC215682500). To inactivate the fluorescence of GFP in cancer cells and background autofluorescence of brain tissues, sections were incubated with 30% H_2_O_2_ (Sigma Aldrich, Cat: 216763500ML) and 0.02 M HCl in PBS under a clamp lamp with 8.5 inch aluminum reflector (Simple Deluxe, Cat: B08MZKQNP4) and a 250W clear bulb (VWR, Cat: 36548001) for 1 hour at 4 °C. The sections were subsequently incubated in the blocking buffer of 5% normal donkey serum (Jackson ImmunoResearch, Cat: 017000121) or 10% normal goat serum (Life Technologies, Cat: 50-062Z), selected to match the host species of secondary antibodies, and 2% (w/v) bovine serum albumin (Thermo Fisher Scientific, Cat: BP9706100) in PBS-T for 1 hour at room temperature. After blocking, sections were incubated in primary antibodies diluted in blocking buffer for overnight at 4 °C, washed in PBS-T six times, and then incubated in 1:500 secondary antibodies and 10 µg/mL DAPI (Thermo Fisher, Cat: D3571) diluted in blocking buffer for 2-3 hours, followed by three washes in PBS-T and three washes in PBS, all at room temperature. Each wash step was 5-10 minutes long. All incubation and wash steps were performed with gentle shaking. Antibodies and corresponding dilutions were detailed below. After the last PBS wash, sections were transferred onto 1.0 mm-thick glass slides (VWR, Cat: 48312003) and allowed to air dry until translucent, mounted with ProLong Gold Diamond antifade mountant (Thermo Fisher, Cat: P36961), and sealed with 0.13- to 0.17 mm-thick glass coverslips (Fisher Scientific, Cat: 125485M). The sealed slides were first cured at room temperature, and placed at −20°C for long-term storage. Images of the slides were taken with a TCS SP5 confocal microscope (Leica Microsystems) or a Ti2-E motorized microscope (Nikon) equipped with Crest X-Light V2 LFOV25 spinning disk confocal (Nikon) and processed as described below (see **Imaging analysis**).

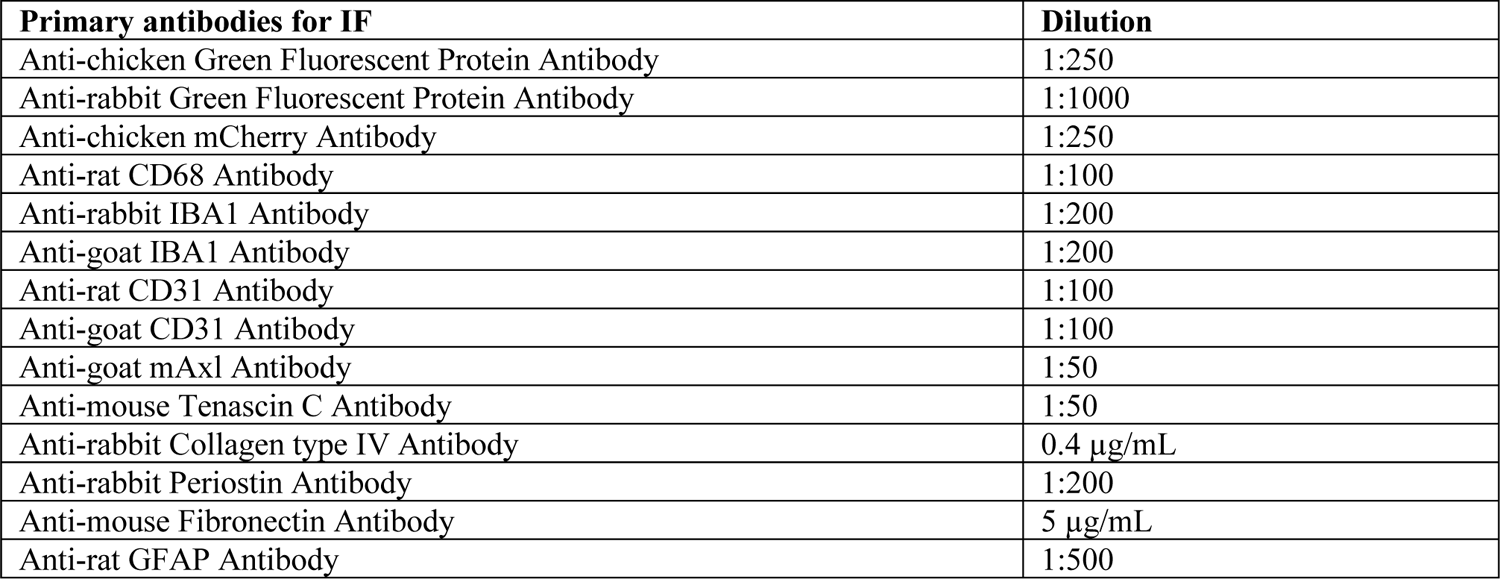

### Whole-mount immunostaining and volume imaging of iDISCO-cleared brain

Immunostaining and imaging of brain hemisphere samples were conducted following the standard iDISCO protocol (idisco.info/idisco-protocol/)^97^ over 4 major steps, 1) pretreatment with methanol (MeOH, Fisher Scientific, Cat: A4524), 2) immunolabeling, 3) tissue clearing, and 4) imaging. Briefly, during 1) pretreatment, harvested tissue samples were dehydrated through serial incubation in 20%, 40%, 60%, 80%, and 100% MeOH/H_2_O solution, each solution for 1 hour and 100% MeOH twice. After incubation with 66% Dichloromethane (DCM, Sigma Aldrich, Cat: 270997-2L)/33% MeOH overnight at room temperature, the dehydrated samples were bleached with 5% H_2_O_2_ in MeOH overnight at 4 °C, and subsequently rehydrated by 1-hour incubation with 80%, 60%, 40%, 20% MeOH/H_2_O and then PBS. Proceeding with 2) immunolabelling, the rehydrated samples were first permeabilized with 20% DMSO (Fisher Scientific, Cat: D128500), 2.3% (w/v) Glycine (Fisher Scientific, Cat: G48212), 0.2% Triton-X in PBS, and then blocked in 10% DMSO, 6% normal donkey serum, 0.2% Triton-X in PBS, each step for 2 days at 37 °C. After blocking, samples were incubated with primary antibodies diluted in 5% DMSO, 3% normal donkey serum in PTwH buffer, consisting of PBS with 0.2% Tween 20 (Sigma Aldrich, Cat: P13791L) and 0.001% Heparin, for 7 days at 37 °C, followed by five washes with PTwH buffer over 1 day at room temperature, and then incubated with secondary antibodies diluted in 3% normal donkey serum in PTwH buffer for 7 days at 37 °C, followed by another five washes with PTwH buffer over 1 day at room temperature. To 3) clear the tissue, immunolabeled samples were dehydrated as in 1) pretreatment by serial incubation in 20%, 40%, 60%, 80%, and 100% MeOH/H_2_O solution, each solution for 1 hour and 100% MeOH twice, and then incubated with 66% DCM/33% MeOH for 3 hours, followed by two 15-minute washes with 100% DCM to remove residual MeOH, with all steps at room temperature. Finally, samples were stored in DiBenzyl Ether (DBE, Sigma, Cat: 108014-1KG) without shaking at 4 °C until imaging. All above steps were performed with shaking except for the final storage. The 5 mL microcentrifuge tubes (Thermo Scientific, Cat: 14568101) used throughout the process were filled with buffer to the top to prevent air from oxidizing the samples. Before 4) imaging, the tubes were inverted a couple times to mix the DBE solution, and cleared samples were transferred to the chamber of a Luxendo MuVi SPIM light sheet microscope (Bruker) to be scanned with a 10X objective, 30% laser power, and in line mode with the resolution of 50 px and light sheet thickness of 3 µm.

| **Primary antibodies for iDISCO** | **Primary antibody dilution** | **Secondary antibodies for iDISCO** | **Secondary antibody dilution** |
| --- | --- | --- | --- |
| Anti-goat CD31 Antibody | 1:100 | Alexa-Fluor 488 Donkey anti-Goat | 1:200 |
| Anti-rabbit IBA1 Antibody | 1:200 | Alexa-Fluor 568 Donkey anti-Rabbit | 1:200 |
| Anti-chicken GFP Antibody | 1:250 | Alexa-Fluor 647 Donkey anti-Chicken | 1:150 |

### Imaging analysis

Confocal microscopy images were minimally processed with FIJI, an implementation of ImageJ (NIH, version 2.3.0/1.53q). Representative images were displayed with linear adjustments of contrast and brightness, using identical adjustment settings where fluorescent signals were quantitatively compared (e.g., AXL and CD68 between MDA231-BrM lesions and HCC1954-BrM lesions in Figure 4A). Figure 1A images showed maximum intensity projection of z-stack images (11 planes, Δz = 0.3 µm), which captured the structure of blood vessels better than single z-plane images. To facilitate visualization of cancer cells immunolabeled by anti-GFP antibody in the 488 nm channel, where brain tissue exhibited a high level of autofluorescence, FIJI functions of Threshold and Analyze Particles were used to mask non-cancer cell regions in cases of high autofluorescence background. No other images were processed by such background removal step. Light sheet microscopy images of a brain hemisphere were registered and stitched by the built-in Image Processor function in Luxendo MuVi SPIM light sheet microscope. The 3D-reconstructed images obtained were adjusted for contrast and brightness in Imaris (Oxford Instruments Group, version 9.5.0). 1 mm^3^ cubic regions containing brain metastases were cropped and displayed as 3D images or videos with optimal rendering.

### Gene overexpression (OE)

To achieve GAS6 OE, HCC1954-BrM cells were transduced with lentivirus expressing *GAS6* under the control of EF1α promoter (pLV[Exp]-BsdEF1A>hGAS6[NM_000820.4], VectorBuilder, Cat: Ecoli(VB900126-3640pet)). As the negative control, cells were transduced with lentivirus carrying a scrambled sequence instead (pLV[Exp]-Bsd-EF1A>ORF_Stuffer, Vector Builder, Cat: Ecoli(VB900122-0480ezn)).

### Gene expression knockdown

#### RNA interference

shRNA-mediated knockdown of *TNC*, *COL4A1*, *GAS6*, and *AXL* expression in HCC1954-BrM cells and MDA231-BrM cells were performed by cloning mir-E-based shRNA sequences targeting these genes into corresponding lentiviral or retroviral vectors listed below to replace their existing shRNA sequences that target the Renilla luciferase gene as negative control^98^. Five 97-mer oligonucleotides (IDT), each containing a different candidate shRNA sequence, were designed per gene using the web-based tool splashRNA (splashrna.mskcc.org)^99^. qRT-PCR analysis was conducted for each engineered cell line to select the top two shRNAs with high knockdown efficiency to target particular gene as shown below.

| shRNA | Plasmid backbone | Vector type | Selection marker | Cells |
| --- | --- | --- | --- | --- |
| sh*TNC* 1 | SGEN<sup>98</sup> | Lentiviral | G 418 (500 μg/mL) (Thermo Fisher, Cat: 10131035) | HCC1954-BrM |
| sh*TNC* 2 | SGEN | Lentiviral | G 418 (500 μg/mL) | HCC1954-BrM |
| sh*COL4A1* 1 | SGEN | Lentiviral | G 418 (500 μg/mL) | HCC1954-BrM |
| sh*COL4A1* 2 | SGEN | Lentiviral | G 418 (500 μg/mL) | HCC1954-BrM |
| sh*AXL* 1 | LENC<sup>98</sup> | Retroviral | FACS sorting for mCherry+ cells | MDA231-BrM,<br>HCC1954-BrM |
| sh*AXL* 2 | LENC | Retroviral | FACS sorting for mCherry+ cells | MDA231-BrM,<br>HCC1954-BrM |
| sh*GAS6* 1 | LEPZ<sup>98</sup> | Retroviral | Puromycin dihydrochloride (5 µg/mL,<br>Sigma Aldrich, Cat: P962010ML) | MDA231-BrM,<br>HCC1954-BrM |
| sh*GAS6* 2 | LEPZ | Retroviral | Puromycin dihydrochloride (5 µg/mL) | MDA231-BrM,<br>HCC1954-BrM |

#### CRISPR interference (CRISPRi)

We used CRISPRi to suppress the expression of *Tnc* and *Col4a1* in MMTV-ErbB2-BrM cells, as shRNA knockdown (KD) of these genes was found to reduce the transcript level only by 50%. MMTV-ErbB2-BrM cells were engineered to express the two components of CRISPRi system – dCas9-KRAB and sgRNAs – via two steps of lentiviral transduction and selection. To first establish cells that stably express dCas9-KRAB, we cloned a lentiviral vector carrying transcription repressor sequence dCas9-KRAB. Specifically, we replaced the VP64-P65-Rta sequence (encoding transcription activator complex) in the lentiviral vector of dCas9-VPR-mCherry fusion protein (for CRISPRa, Addgene #102245) with the KRAB-MECP2 sequence from the transient expression vector of dCas9-KRAB-MECP2 (for CRISPRi, Addgene #110821)^100^. After lentiviral transduction, we isolated Cherry+ cells by FACS sorting, which were then introduced with specific sgRNAs targeting *Tnc* or *Col4a1* as the second step as follows. The sgRNAs listed below, including the one targeting Gal4 DNA binding domain that was designed as a negative control, were cloned into the lentiGuide-Hygro-mTagBFP2 vector (Addgene #99374) as previously described^101^. Specifically, in synthesizing the oligonucleotides (IDT) to clone sgRNAs, 5’CACC and 5’AAAC overhangs were added to sense and antisense oligonucleotides, respectively, to create cohesive ends in annealing. The annealed oligonucleotides were ligated into the backbone of lentiGuide-Hygro-mTagBFP2 vector, obtained by BsmBI-v2 (New England Biolabs, Cat: R07395) digestion. The mCherry+ tagBFP+ cells were FACS-sorted after the second round of lentiviral transduction, and tested for the efficiency in knocking down *Tnc* or *Col4a1* expression by qRT-PCR.

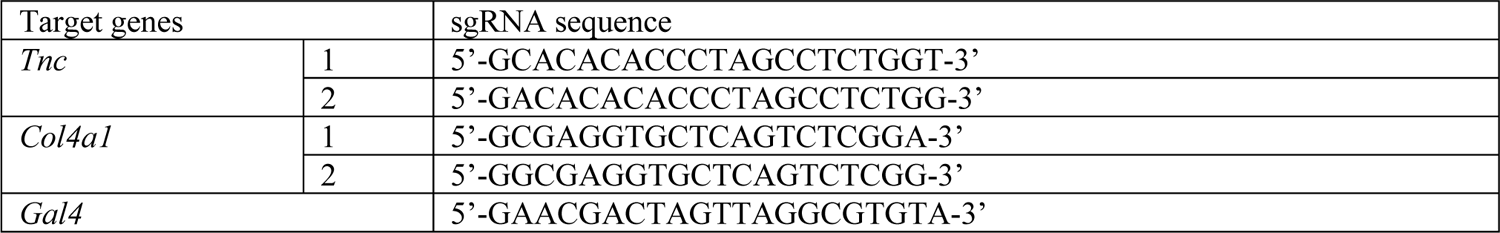

### RNA isolation and bulk gene expression assays

#### RNA extraction

Cells grown to approximately 80% confluence in a 10 cm petri dish (Fisher Scientific, Cat: 08772E) were collected as a cell pellet for total RNA extraction. Cells were lysed through QIAshredder columns (Qiagen, Cat: 79656) in Buffer RLT from QIAgen RNeasy Mini Kit (QIAgen, Cat: 74106) supplemented with 1:100 β-Mercaptoethanol (Sigma Aldrich, Cat: M3148100ML). After cell lysis, RNA isolation was performed with the RNeasy Mini Kit. Total extracted RNA was quantified using a NanoDrop (Fisher Scientific, Cat: 2741002PM22). Reverse transcription was performed using the Transcriptor First Strand cDNA Synthesis Kit (Roche, Cat: 04897030001).

#### qRT-PCR

For quantitative real-time polymerase chain reaction (qRT-PCR), 1 µg of total RNA was used to synthesize cDNA. Amplification of targets from cDNA was performed with the TaqMan Universal PCR Master Mix (ThermoFisher, Cat: 4304437) and TaqMan probes in a ViiA 7 Real-Time PCR System (ThermoFisher) via 40 cycles of PCR, each consisting of 10 seconds at 95 °C (denaturation) and 30 seconds at 60 °C (annealing/polymerization). Relative gene expression was calculated by the canonical 2**^−^**^ΔΔ*C*^_T_ method^102^, using housekeeping genes *ACTB* or *GAPDH* as control.

#### Flura-seq

As summarized in Figure S6A, *Flura-seq* was conducted as previously described^103, 104^ with minor modifications. Briefly, MDA231-BrM cells and HCC1954-BrM cells were transduced with lentivirus carrying PGK-UPRT-T2A-CD. For each BrM model, 1.0 × 10^5^ cells resuspended in 100 µL PBS supplemented with 2% FBS were inoculated into the mice by intracardiac injection as described above (see **Animal Studies**). 30 days post-inoculation, mice were intraperitoneally injected with 250 mg/kg 5-fluorocytosine (5-FC) (Sigma Aldrich, Cat: F71291G), and after 12 hours, mouse brains were harvested and homogenized. mRNA was extracted from homogenized cell lysate by 4 × 250 µL/brain Oligo (dT)_25_ magnetic beads (New England Biolabs, Cat: S1419S), and immunoprecipitated (IP) with 2.5 µg/brain anti-BrdU antibody (Abcam, Cat: ab6326). cDNA libraries were constructed using the immunoprecipitated mRNA and SMARTer PCR cDNA Synthesis Kit (Clontech, Cat: 634926), and sequenced on a HiSeq 2500 system (Illumina, Integrated Genomics Core, MSKCC).

#### RNA-seq

RNA-seq samples were processed and sequenced in the Integrated Genomics Core, MSKCC. The quality and quantity of RNA samples were determined using BioAnalyzer 2100 (Agilent). cDNA libraries were constructed using either TruSeq RNA Sample Prep Kit v2 (Illumina, Cat: RS-122-2001) or NEBNext Ultra RNA Library Prep Kit (New England Biolabs, Cat: E7530S) and sequenced on a HiSeq 2500 system (Illumina) at a depth of 25-50 million reads per sample.

### Bulk gene expression data analysis

#### Expression signatures of brain tropism

RNA-seq profiles of HCC1954 and MMTV-ErbB2, including parental cells and BrM derivatives, were generated in this paper. Microarray gene expression data of MDA231 parental cells and BrM derivatives were obtained from previous breast cancer dataset^3^. Raw sequencing reads in FASTQ format were mapped by STAR RNA-seq aligner^105, 106^ (version 2.5.3a), using human genome reference GRCh38 and mouse genome reference GRCm38 from GENCODE (www.gencodegenes.org/) for RNA-seq samples of human cancer cells and mouse cancer cells, respectively. Uniquely mapped reads were assigned to annotated genes by HTSeq (version 0.6.1p1) with default settings^107^ to measure read counts. The counts were normalized by library size to be subject to differential gene expression analysis, both performed with the DESeq2 package^108^ (Bioconductor version 3.4) in RStudio (version 1.0.153) implementing R (3.4.1). A gene was defined as differentially expressed if its 1) absolute value of log2 fold change in normalized counts was greater or equal to 2, 2) adjusted P value was below 0.05, and 3) mean read counts was larger than 10. Flura-seq data was analyzed following the same procedure with one additional step preceding alignment. As the data were derived from xenograft mouse model samples containing a mixture of reads from human cancer cells and mouse stromal cells, Xenome^109^ (version 1.0.1) was run as the first step to classify the reads as belonging to either human genome or mouse genome using the default value (25) of parameter *k*-mer size. The web-based tool DAVID^77^ (Database for Annotation, Visualization and Integrated Discovery, version 6.8) was employed to identify over-represented gene ontology (GO) terms in differentially expressed genes (DEGs).

#### Analysis of MetMap dataset

Bulk RNA-seq profiles of brain metastases with multiplexed breast cancer cell line composition were obtained from the MetMap dataset, and the differential expression analysis that properly accounted for cancer cell composition variability was conducted as previously described^27^. The percentage of cells belonging to HER2BC versus TNBC cell lines in a sample were computed. To assess the statistical significance of the *in vivo* upregulation of genes encoding ECM components in brain metastases in relation to the HER2BC subtype, linear regression analysis was performed on the log2 fold changes in gene expression with respect to the fraction of HER2BC cells. Wilcoxon rank sum test was performed on the log2 fold changes in gene expression with respect to the dominance of HER2BC cells. P values were adjusted by BH multiple hypothesis correction. Data were plotted using ggplot2 in RStudio (version 1.2.5019).

#### Query of GAS6 and AXL levels in breast cancer cell lines in DepMap database

For all human breast cancer cell lines available in the DepMap database, consistent subtype annotation, including HER2BC, TNBC, luminal A, and luminal B, were confirmed by cross-validating Ref.^20^, DepMap, and breast cancer cell line classification resources^21^. DepMap transcriptomics data were originally processed through STAR alignment, RSEM quantification^110^, and TPM (transcripts per million) normalization. The normalized transcript levels (log2(TPM + 1) values) were obtained from the DepMap online portal (depmap.org/portal/). Proteomics data were measured by mass spectrometry^111^, and the relative normalized protein levels published by Ref.^111^ were obtained from the DepMap online portal as well. Correlation between AXL and GAS6 was assessed by subtype-specific Spearman’s correlation analysis. Transcript and protein levels were compared across subtypes using a Wilcoxon signed-rank test. All analysis was performed using R (version 4.2.0, www.r-project.org) and Bioconductor (version 3.15).

#### scRNA-seq data collection

HCC1954-BrM cells and MDA231-BrM cells were transduced with lentivirus expressing sLP-mCherry-P2A-eGFP under the control of EF1α promoter (plasmid constructed by Tammela Lab, MSKCC). As illustrated in Figure 2C, athymic mice were intracardially inoculated with the engineered MDA231-BrM cells or HCC1954-BrM cells (1.0 × 10^5^ cells/mouse). A mouse brain harboring MDA231-BrM metastases and one harboring HCC1954-BrM metastases were harvested 4 weeks post-inoculation as described above (see **Animal studies**), and both processed through 1) enzymatic tissue dissociation using the Adult Brain Dissociation Kit, mouse and rat (Miltenyi Biotec, Cat: 130-107-677), 2) removal of myelin using Myelin Removal Beads II, human, mouse, rat (Miltenyi Biotec, Cat: 130-096-733), and 3) removal of debris and dead cells using the Dead Cell Removal Kit (Miltenyi Biotec, Cat: 130-090-101). The brain of an age-matched uninoculated mouse was harvested and processed in parallel to serve as the negative control for downstream FACS sorting. The first enzymatic tissue dissociation step occurred at 37 °C, and all following steps were performed at 4 °C.

The resulting single-cell suspension of each brain metastasis sample was incubated with an anti-human Hashtag antibody (binding to human cancer cells) and an anti-mouse Hashtag antibody (binding to mouse cells), both conjugated to unique DNA barcodes as listed below, 1:100 human TruStain FcX (Fc Receptor Blocking Solution) (BioLegend, Cat: 422301) and 1:100 TruStain FcX (anti-mouse CD16/32) antibody (BioLegend, Cat: 101319) to prevent unspecific binding of Hashtag antibodies, and 40 ng/mL CellTrace Calcein Violet, AM, for 405 nm (Thermo Fisher Scientific, Cat: C34858) and 1:20 APC Annexin V BioLegend, Cat: 649020) to allow filtering residual debris and apoptotic cells during FACS sorting. The Calcein Violet+ Annexin V− live cells of each sample were FACS-sorted into 3 groups, i.e., the labeling GFP+ mCherry+ cancer cells, labeled GFP− mCherry+ TME cells, and rest unlabeled GFP− mCherry− cells. The single-cell suspension of uninoculated mouse brain, which only contained GFP− mCherry− cells, was used to set the gates for GFP+ or mCherry+ cells.

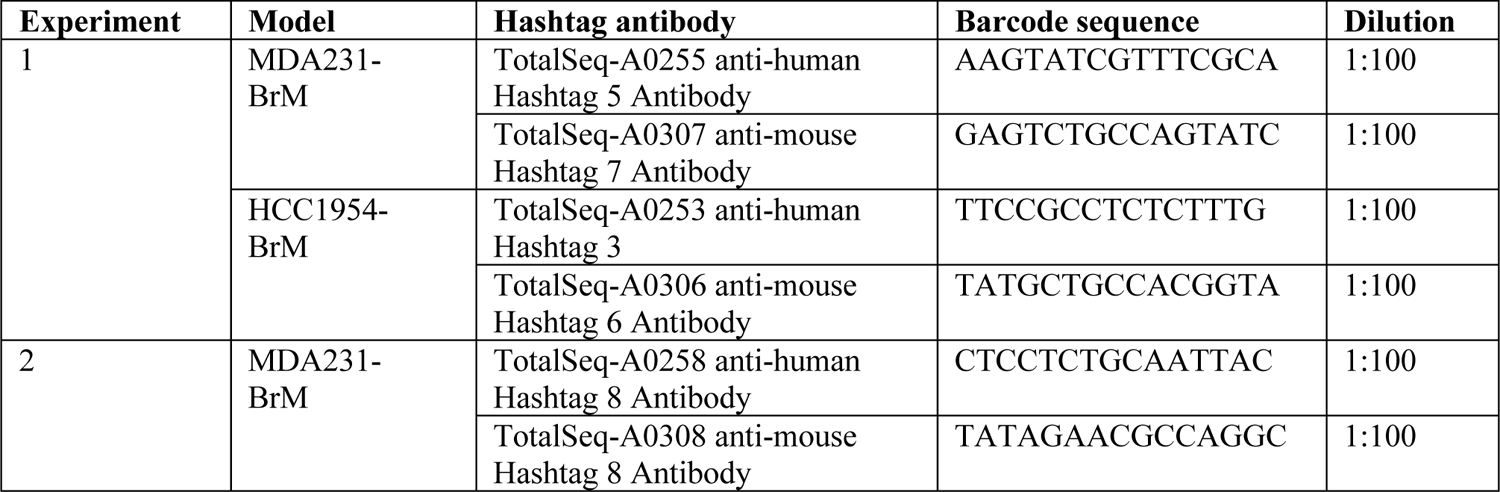
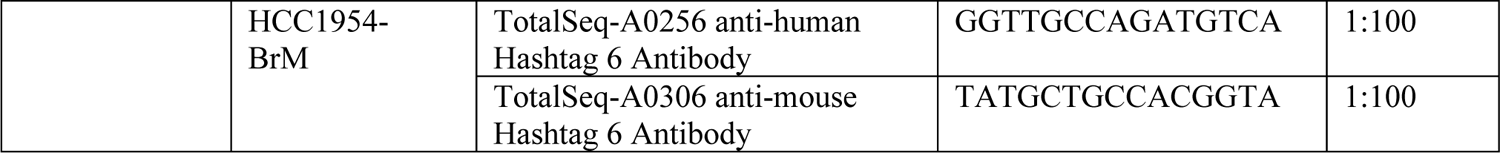

The sorted cells from MDA231-BrM and HCC1954-BrM samples were then pooled per FACS-sorted group for single-cell droplet encapsulation and sequencing, and subsequently assigned to their original sample source according to the sequencing reads of different Hashtag sequences used (see *<u>Pre-processing of scRNA-seq data</u>* in **scRNA-seq data analysis**). Two independent scRNA-seq experiments were conducted following similar steps. In experiment 2, 5 μg/mL Actinomycin D (Sigma, Cat: A1410), 10 μM Triptolide (Sigma, Cat: T3652), and 27.1 μg/mL Anisomycin (Sigma, Cat: A9789) were added to the PBS used for perfusing mice and soaking the harvested brains and to the enzymatic dissociation buffer as well, in order to inhibit *ex vivo* transcription and translation as previously reported^45^.

#### scRNA-seq data analysis

Data from both experiments (1, 2) were combined to be pre-processed (i.e., filtering ambient RNA and non-cell droplets, normalizing and log-transforming single-cell UMI counts, calling cell types, computing signature scores) together before being separated for downstream analysis following the same procedure, presented in Figures 2, S2, S3 (experiments 1, 2), 3 (experiment 1) and S4 (experiment 2). All our analysis results were consistent between two scRNA-seq experiments, except that in the second one, pro-inflammatory cytokine gene transcripts were diminished likely due to the transcription and translation inhibitors added^45^.

#### Pre-processing of scRNA-seq data

FASTQ files of sequenced samples were individually processed using the SEQC pipeline^54^ (version 0.2.1) with default parameters for the 10x single-cell 3’ library. Samples from GFP+ cancer cells and from GFP− non-cancer cells in the brain were mapped to the hg38 human and mm38 mouse genome references, respectively. The SEQC pipeline performed read alignment, multi-mapping read resolution, and cell barcode and UMI correction to generate a raw UMI-based (cell × gene) count matrix per sample. Following SEQC, we ran multiple complementary quality control (QC) algorithms in parallel, and used the Scanpy software^112^ (version 1.6.0) in Python (version 3.8.5) to incorporate their results to identify high-quality, background-removed single-cell transcriptome from the count matrices. The algorithms used were: 1) CellBender^113^ (version 0.2.0) removed counts due to ambient RNA molecules and random barcode swapping from the (raw) SEQC output count matrices, using the remove-background command with the parameters of expected-cells = 10000, total-droplets-included = 50000, fpr = 0.01 (default value), and epochs = 150 (default value). 2) DropletUtils^114^ (version 1.10.3) distinguished the droplets containing cells from those containing ambient RNA by the function of emptyDrops on the (raw) SEQC output count matrices with the parameters of retain = Inf, test.ambient = True, lower = 50, niters = 50000. 3) An in-house method called SHARP (github.com/hisplan/sharp, version 0.2.1) assigned a cell as labeled by a particular Hashtag oligo and thereby belonging to the specific mouse (bearing either MDA231-BrM metastases or HCC1954-BrM metastases) barcoded with this oligo or as a doublet or low-quality droplet. We subsequently uploaded the CellBender-processed count matrices, subset to droplets that were called to be real cells by both emptyDrops (FDR ≤ 0.01) and SHARP, and computed additional metrics on the subset matrices to eliminate residual droplets that were likely to be empty droplets, doublets or apoptotic cells. We used Scanpy’s pp.normalize_total function to normalize the count matrices by library size (i.e., total number of UMIs after CellBender filtering of background counts). Using the library size-normalized count matrices (base = 2, pseudo count = 1), we performed principal component analysis (PCA) using the top 5000 highly variable genes (HVGs) that were robustly detected (in at least 10 cells), excluding mitochondrial or ribosomal genes, and ran PhenoGraph^115^ (version 1.5.6, *k* = 30) to cluster the droplets. To exclude apoptotic cells, we filtered out droplets with a high fraction of mitochondrial gene transcripts (mtRNA% ≥ 0.2), indicative of cellular stress. To remove droplets of low library complexity (i.e., with few unique genes), we followed the SEQC^54^ approach of fitting a linear model to the relationship between the number of genes and the number of UMIs across droplets, and discarded droplets with lower than expected number of genes (residuals > 0.15, default value in SEQC). To clean out any remaining lower-quality droplets, we examined the gene-gene expression covariance of each PhenoGraph cluster computed on its top 500 highly expressed genes and removed cells constituting the PhenoGraph clusters that did not exhibit apparent covariance structures. After the above filtering steps, we ran DoubletDetection (version 2.5.2, github.com/dpeerlab/DoubletDetection) to infer and remove droplets that may contain more than one cell. The above pre-processing steps yielded:

| Experiment | Cells | FACS group | Source | Number of cells | Number of genes | Median of UMIs/cell | Median of genes/cell |
| --- | --- | --- | --- | --- | --- | --- | --- |
| 1 | Cancer | GFP+ mCherry+ | TNBC | 5659 | 19284 | 15185 | 3866 |
|  |  |  | HER2BC | 2025 | 18610 | 20773 | 4356 |
|  | Non-cancer | GFP- mCherry- (Unlabeled) | TNBC | 2303 | 16992 | 3929 | 1704 |
|  |  |  | HER2BC | 8570 | 18940 | 4232 | 1783 |
|  |  | GFP- mCherry+ (Labeled) | TNBC | 5691 | 16820 | 4756 | 1749 |
|  |  |  | HER2BC | 5187 | 17492 | 6816 | 2093 |
| 2 | Cancer | GFP+ mCherry+ | TNBC | 2444 | 18045 | 11062 | 3499 |
|  |  |  | HER2BC | 3054 | 19016 | 16792 | 4157 |
|  | Non-cancer | GFP- mCherry- (Unlabeled) | TNBC | 1790 | 17963 | 6443 | 2440 |
|  |  |  | HER2BC | 4644 | 18980 | 7335 | 2623 |
|  |  | GFP- mCherry+ (Labeled) | TNBC | 1054 | 17425 | 13623 | 3490 |
|  |  |  | HER2BC | 2418 | 18518 | 14167 | 3706 |

#### Basic analysis of single-cell transcriptome

After constructing a count matrix as described in *<u>Pre-processing of scRNA-seq data</u>*, we concatenated the original (un-normalized) CellBender-processed count matrices of both experiments (1, 2) for filtered GFP+ cancer cells (GFP+ mCherry+) and GFP− non-cancer cells (GFP− mCherry−/unlabeled and GFP− mCherry+/labeled), respectively, and used scran^116^ (version 1.14.6) to normalize the concatenated count matrices. We chose scran instead of the simpler library size normalization performed for pre-processing, because scran is designed to better recapitulate the endogenous variability in library size among a heterogeneous population of cells, and has been verified to be a top-performing normalization method by multiple independent comparisons^117^. After scran normalization, we log-transformed the count matrices (base = 2, pseudo count = 1), and performed principal component analysis (PCA) using the top 5000 highly variable genes (HVGs) that were robustly detected (in at least 10 cells), excluding mitochondrial or ribosomal genes. The top principal components (PCs) selected according to the knee point of total variance explained (as listed below) were used to construct *k*-nearest neighbor (*k*NN) graph (n_neighbors = 15) to generate uniform manifold approximation and projection (UMAP) layouts, and to cluster cells by PhenoGraph^115^ (version 1.5.6, *k* = 30). The PhenoGraph clusters of GFP− non-cancer cells were proceeded with for cell-type annotation.

| Cells | Number of PCs | Variance explained |
| --- | --- | --- |
| Cancer | 56 | 0.43 |
| Non-cancer | 42 | 0.56 |

#### Cell-type annotation of non-cancer cells

We annotated the cell types of GFP− non-cancer cells using a reference single-cell transcriptome atlas of the mouse nervous system^46^ (hereinafter referred to as “atlas”, mousebrain.org) and additional immune cell type markers that complement the atlas (see Figure S2, Table S1). The atlas determined a hierarchical classification of cells in the nervous system. At the most refined level of classification, it identified 265 cell types, and provided 1) cluster-averaged expression, 2) six marker genes, and 3) associated metadata including the upper-level taxonomy annotation of the clusters. The additional immune cell type markers were collected from commonly used scRNA-seq cell type markers and included genes for calling B cells (BC), neutrophils (NEUT), bone marrow-derived macrophages (BMDM), and NK cells (NK) – infiltrated immune cells absent in the atlas database of healthy brain tissue. See Table S1 for cell type marker genes and metadata.

We first mapped the query PhenoGraph^115^ clusters (see *<u>Basic analysis of single-cell transcriptome</u>* in **scRNA-seq data analysis**) to their respective reference cell type by evaluating two metrics – 1) cellular detection score of marker genes and 2) correlation of global gene expression levels. 1) Cellular detection score was quantified as described previously^54^. For a given cell *c* and marker gene *k*:

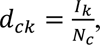

where a cell was scored 1 (*I*_k_ = 1) if it contains a non-zero UMI count of the marker gene and 0 otherwise (*I* = 0), and corrected by the detection rate of the cell, defined as the fraction 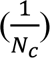 of total number of genes detected in it (*N*_c_). The corrected scores (*d*_ck_) were subsequently averaged across marker genes of a reference cluster and all cells from certain query PhenoGraph cluster to generate the query cluster-level cellular detection score for certain group of marker genes as:

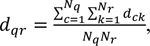

where *N*_q_ is the total number of cells in the query cluster *q*, and *N*_r_ total number of marker genes in the reference cluster *r*. 2) Correlation in cluster-level global expression profiles. Specifically, we summarized the gene expression of query clusters by averaging single-cell normalized UMI counts across cells per query cluster. We obtained the cluster-level normalized UMI counts of 265 reference clusters from the atlas database. We then computed the correlation in the log-transformed cluster-averaged count matrices between query PhenoGraph clusters and 265 reference atlas clusters on 5036 robustly detected genes. These genes were selected from genes with ≥ 1 average normalized UMI counts per query cluster, excluding abundant mitochondrial and ribosomal genes. The selection was performed to ensure that the correlation computed was representative of the expression patterns of all query clusters, and not overwhelmed by individual clusters with higher number of cells (e.g., macrophages) or genes ubiquitously expressed at high or low levels across clusters. Correlation with infiltrated immune cell types was not computed, as the atlas does not contain transcriptome data of these cell types. For each query cluster of non-cancer cells, we noted that both metrics had one or several high-scored reference clusters which were well separated from the rest, and that the top-scored reference cluster(s) were largely identical or overlapping between metrics, indicative of consistent cell type calling by either specific marker genes or global gene expression (if available). We manually examined the common (of both metrics) top-scored reference clusters for each query cluster, especially in the cases of more than one reference clusters and in most of such cases, the reference clusters shared the same upper-level taxonomy (e.g., MGL1, MGL2, and MGL3 clusters in the atlas corresponding to different states of the same taxonomy of microglia, MG), which we used to annotate the cell population. In the remaining rare cases where multi-mapping arose from certain query cluster containing cells of closely related but different cell types (e.g., MG, BMDM, and border associated macrophages, BAM, all belong to macrophages), we repeated the PCA and PhenoGraph clustering for this query cluster specifically to obtain more refined grouping and re-ran the above cell type annotation steps of re-grouped query clusters. After finalizing the annotation through an iterative optimization process, we re-computed and plotted the two metrics restricted to identified populations and relevant reference clusters to generate the plots in Figures 2E, 2F, and S2.

#### Inferring tissue dissociation-associated ex vivo activation and cell cycling status in macrophages

As noted above (see *<u>Cell-type annotation of non-cancer cells</u>*), macrophages include microglia (MG), bone marrow-derived macrophages (BMDM), and border associated macrophages (BAM). To infer the cell cycling status of macrophages and evaluate the impact of adding transcription and translation inhibitors^45^ during tissue dissociation in experiment 2, we utilized the score_genes function of Scanpy^112^ to compute on all macrophages the signature scores of 1) genes associated with S phase or G2/M phase^118^ and 2) genes whose expression can be induced *ex vivo* by the enzymatic dissociation process^45^, respectively (both listed in Table S2). The Scanpy^112^ gene signature score is calculated as the average expression of a set of genes, subtracted with the average expression of a reference set of genes that were randomly sampled to match the expression distribution of the given gene set. To ensure that the score values of the cells from two experiments were directly comparable to each other, signature scores were computed on both experiments together by running the score_genes function once on their concatenated matrices rather than twice on separate matrices. Two PhenoGraph clusters of microglia displayed markedly higher scores of S phase and G2/M phase genes, and were inferred to be cycling (Figure S3C) (and the majority of microglia as non-cycling, in G0 or G1 phase). The substantially lower scores of tissue dissociation-associated *ex vivo* activation (Figure 2G) in the macrophage cells from experiment 2 validated the reported effects of including transcription and translation inhibitors^45^.

#### Differential sample abundance visualization and analysis

We adapted the Milo algorithm^47^ (version 0.1.0) built upon the generalized linear model (GLM) to analyze the differential abundance in sub-populations between conditions. Milo expects replicates in its input but is exceedingly sensitive to batch effects and even minor imperfections in data integration. Thus, we chose to analyze experiment 1 and 2 separately, which required a change in Milo’s test statistic, and looked for correspondence in Milo’s results across the two experiments. We aimed to compare the subpopulation frequencies in MDA231-BrM TNBC- or HCC1954-BrM HER2BC-labeled (GFP− mCherry+) cells in reference to unlabeled (GFP− mCherry−) cells. We isolated the log-transformed, normalized UMI counts of the cells of interest (i.e., either all non-cancer cells or non-cycling microglia only) from each experiment to construct a *k*NN graph (*k* = 15) and generate UMAP embeddings, using the following indicated number of PCs computed as described above (see *<u>Basic analysis of single-cell transcriptome</u>* in **scRNA-seq data analysis**). As described in *<u>Identifying gene programs associated with homeostasis-to-DAM transition</u>*, the same number of PCs were also selected to compute diffusion components, using the Palantir algorithm^55^ (version 1.1) that runs diffusion maps with an adaptive anisotropic kernel. To intuitively visualize the differential abundance in the phenotypic space, we used the Seaborn kernel density estimate (KDE) plot (with 7 contour levels) to show the distribution of each group of non-cycling microglia cells in the UMAP embedding computed on all 4 groups of cells (i.e., TNBC-labeled, TNBC-unlabeled, HER2BC-labeled, HER2BC-unlabeled) together.

| Experiment | Cells | Number of PCs | Variance explained |
| --- | --- | --- | --- |
| 1 | All non-cancer cells | 43 | 0.51 |
|  | Non-cycling microglia | 98 | 0.24 |
| 2 | All non-cancer cells | 40 | 0.61 |
|  | Non-cycling microglia | 128 | 0.41 |

We followed the steps of the Milo algorithm^47^ to select index cells, using a graph sampling strategy previously devised by Palantir^55^, and to construct partially overlapping neighborhoods defined on the index cells. To increase statistical power, since both TNBC-unlabeled and HER2BC-unlabeled groups comprised cells distal to tumor lesions in the brains of athymic mice, we assumed these two groups as the replicates of unlabeled cells. Let μ_ns_ denote the mean counts of cells of group *S* in neighborhood *n, M_s_* the total number of cells of group *S*, and *I_sTNBC-labeled_* and *I_sHER2BC-labeled_* the indicator variables of whether group *s* is TNBC-labeled (*I_sTNBC-labeled_* = 1 for the TNBC-labeled group, and 0 otherwise) or HER2BC-labeled (similarly, *I_sHER2BC-labeled_* = 1 for the HER2BC-labeled group, and 0 otherwise), following the Milo framework, we modeled the cell counts from certain neighborhood *n* as

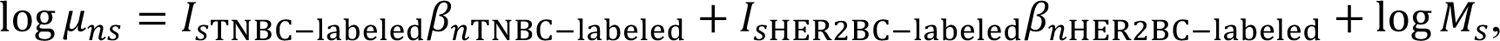

where β*_nTNBC-labeled_* and β*_sHER2BC-labeled_* corresponded to the regression coefficients by which the effects of originating from the TNBC-labeled or HER2BC-labeled group were mediated for neighborhood *n*. We deviated from Milo in the statistical test applied to identify significantly differential abundances from each experimental condition per neighborhood. Instead of the quasi-likelihood F-test selected by Milo, we utilized alternative log-likelihood ratio test functions in EdgeR package^119^ (version 3.32.1) best fit for 1-2 replicates per condition, smaller cell counts, and larger dispersion in the cell counts. To evaluate the variability in cell counts in replicate samples for each neighborhood, we ran the EdgeR^119^ function estimateDisp with default parameters, which maximized the negative binomial likelihood to estimate the dispersion in cell counts across phenotypic neighborhoods. To obtain β*_nTNBC-labeled_* and β*_sHER2BC-labeled_* values as the log fold change (logFC) in labeled cell abundance, we ran glmFit to fit a negative binomial generalized log-linear model to the cell counts of each neighborhood. To test whether TNBC-labeled or HER2BC-labeled sample abundance differed from unlabeled sample abundance, we ran glmLRT to conduct log-likelihood ratio test on whether β*_nTNBC-labeled_* or β*_sHER2BC-labeled_* differed from 0. Lastly, we applied the Milo function graphSpatialFDR using default parameter values to correct the glmLRT-derived P values for multiple testing while accounting for the non-independence of overlapping neighborhoods. The corrected P values and logFC values were used for visualization (Figures 3B, 3E, S3B, and S4D) and downstream analysis. It should be noted that, compared to the quasi-likelihood F-test which accounts for uncertainty in dispersion estimates to obtain more conservative and rigorous type I error rate control, the log-likelihood ratio test runs a higher risk of generating false positives^119^. Aware of this potential caveat, we based the majority of our downstream analysis on tracking expression patterns along with continuous changes in logFC values without thresholding on corrected P values, and utilized the scRNA-seq analysis to generate hypotheses which we tested experimentally.

#### Differential gene expression in tumor-associated microglia

To detect genes characteristic of tumor-associated microglia in both subtypes of breast cancer in experiment 1, we first identified two groups of phenotypic neighborhoods in non-cycling microglia that were enriched in or depleted of tumor-associated microglia, respectively, and ran MAST^120^ (version 1.16.0) to call genes differentially expressed between these two groups of microglia. As the first step, neighborhoods that were connected in *k*NN graph (i.e., sharing at least one common cell) and exhibited concordant significant differential abundance in TNBC-labeled and HER2BC-labeled cells (i.e., corresponding logFC values being both positive or both negative, BH-adjusted P values ≤ 0.1) were merged into two groups of neighborhoods – one enriched in (positive logFC values) and one depleted of (negative logFC values) labeled non-cycling microglia from both brain metastasis models. By comparing between the transcriptome of cells in these two groups using MAST, 838 DEGs with absolute log2 fold change ≥ 0.322 (= log2(1.25)) and Benjamini-Hochberg (BH)-adjusted P value ≤ 0.01 were defined to be differentially expressed in tumor-associated microglia in reference to unassociated homeostatic microglia. Given fewer non-cycling microglia captured in experiment 2 (1737 cells, compared to 11407 cells from experiment 1), to enhance robustness in calling DEGs, we compared cells from connected phenotypic neighborhoods that displayed concordant significant enrichment in TNBC-labeled and HER2BC-labeled cells (i.e., corresponding logFC values being both positive, BH-adjusted P values ≤ 0.1) to all remaining cells, which allowed pooling more cells to be used as control. Running MAST with the same thresholds identified 546 DEGs between these two groups of cells, 501 of which (92%) overlapped with the list of 838 DEGs from experiment 1. As shown in Figure S4C, the logFC values of experiment 2 spanned a narrower dynamical range compared to experiment 1, possibly due to the limited total number of cells, which affected statistical power, and the less restricted selection of unlabeled control microglia that was performed to counteract cell number limitation. As evaluated by linear regression performed with the scikit-learn library (version 0.24.1), despite differences in amplitude, the logFC values of all 11379 genes tested by MAST (which dropped inestimable genes) were well correlated between experiments 1 and 2 (*R*^2^ = 0.96, see Table S3 for MAST results), indicating consistent metastasis-induced changes in the global transcriptome of microglia in both experiments.

#### Identifying gene programs associated with homeostasis-to-DAM transition

Considering potentially higher sensitivity in calling DEGs in experiment 1, we used its resulting larger set of 838 DEGs to identify gene programs associated with homeostasis-to-DAM transition by pre-ranked gene set enrichment analysis of these DEGs (Table S3). The analysis was run using the GSEA software^121, 122^ (version 4.0.3) with the default of 1000 permutations. DEGs were pre-ranked according to their Spearman correlation coefficients, calculated with the SciPy package (version 1.4.1) in python, between the diffusion component 1 (DC1) values and neighborhood-level gene expression of all index cells of phenotypic neighborhoods. As noted above, DC1 values were computed by Palantir^55^ on the 98 PCs of non-cycling microglia from experiment 1 (see *<u>Differential sample abundance</u>* in **scRNA-seq data analysis**). Neighborhood-level expression of a given gene and index cell was calculated as the mean log-transformed, normalized UMI counts of the gene among cells in the corresponding neighborhood. GSEA results obtained using a comprehensive collection of reference gene sets, including curated gene programs (organized in Table S2) and mouse Hallmark (MH) and Biological Process (BP) subset of ontology gene sets (M5)^121, 123, 124^ displayed multiple positively enriched oxidative phosphorylation-related BP gene sets that largely overlapped with each other and with the oxidative phosphorylation gene set of Hallmark as well. To reduce redundancy and provide more informative graphical presentation, Figure 3D summarized the results of GSEA run without BP gene sets (see Tables S4, 5 for both sets of GSEA results, run with or without BP gene sets). Among the gene sets shown, oxidative phosphorylation and glycolysis gene sets were obtained from the Hallmark database^123^; DAM signature (88 genes)^48^; axon tract-associated microglia (ATM) signature and CNS pathology-shared microglia responses (i.e., a core, common set of 12 genes upregulated in ATM, DAM, and LPC-induced injury responsive microglia or IRM in abbreviation)^50^; TGF-β-dependent genes, determined specifically in microglia either *in vitro* or *in vivo*^56^; genes dependent on LRRC33 (a membrane-anchored carrier and activator of latent TGF-β required for microglia homeostasis)^57^; homeostasis signatures^49, 125^, all provided in Table S2.

#### Visualizing gene expression patterns

We computed the signature scores for homeostasis and DAM-related gene programs (all listed in Table S2) on individual non-cycling microglia cells from both experiments, using the score_genes function of Scanpy^112^ as described above (see *<u>Inferring tissue dissociation-associated ex vivo activation and cell cycling status in macrophages</u>*<u>). We then calculated the neighborhood-level signature scores in particular experiment as the mean scores of cells in a neighborhood, similar to that of the neighborhood-level gene expression of individual DEGs (see *Identifying gene programs associated with homeostasis-to-DAM transition* in **scRNA-seq data analysis**). To fit the trends of neighborhood-level signature scores and gene expression along the DC1 values of phenotypic neighborhood index cells (see *Differential sample abundance analysis* for sampling index cells), we used Palantir</u>^55^<u>, which employs the generalized additive model (GAM)</u>^126^ <u>to derive robust estimate of the nonlinear expression trends and estimate the standard error of prediction</u>^126^<u>. As a convenient way of implementing the trend fit, we ran the compute_gene_trends function in Palantir</u>^55^<u>, using the neighborhood-level expression and DC1 values of index cells (instead of the default single-cell expression and pseudotime) as the input of the function. As visualized by the plot_gene_trend_heatmaps function of Palantir</u>^55^<u>, the GAM-predicted expression trends were presented as heatmaps, with the values z-normalized across the index cells per gene or gene set. To detect genes whose expression trends followed the differential abundance in TNBC-labeled microglia along DC1, we clustered the DEGs by the cluster_gene_trends function of Palantir</u>^55^<u>, which z-normalizes the gene expression trends (as performed for visualization) to bring the DEGs to the same scale, and performs PhenoGraph</u>^115^ <u>clustering on the z-normalized trends, using a high value of *k =* 150 to avoid over-fragmented clustering (see</u> Table S3 <u>for full clustering results). We identified the cluster 4 genes that contained *Tnf*, *Ccl4*, and *Il1b*, whose transcripts were enriched in TNBC-associated microglia and known to be potently upregulated in the stage 1 DAM of</u> Alzheimer’s disease^63^. The gene trend clustering was performed only for experiment 1, as the aforementioned genes that encode proinflammatory cytokines were found to be depleted in experiment 2 possibly due to the transcription and translation inhibitors added^45^. All other steps were conducted for both experiments 1 and 2, which replicated the transition from homeostasis to differential stages of DAM in metastasis-associated microglia.

## QUANTIFICATION AND STATISTICAL ANALYSIS

### Statistical analysis for non-sequencing data

Sample sizes were indicated in figures or figure legends. Statistic tests were performed in the Prism software (GraphPad, version 8.4.3). P values were computed by two-tailed Mann-Whitney U test unless otherwise noted. Box-whisker plots were shown to summarize the values of minimum, lower quartile, median, upper quartile, and maximum. Web-based survival analysis tool *km-plotter* (www.kmplot.com) was used to assess the correlation between Affymetrix gene expression profiles and relapse-free survival in the clinical samples of HER2BC breast cancer patients (n = 695). For each queried gene, *km-plotter* classified all samples into two cohorts using three pre-defined quantiles (i.e., median, upper and lower quartiles) as the cutoff expression value, computed Cox (proportional hazards) regression for each cutoff value, and selected the quantile that yielded the highest significance (lowest *P* values) as the best cutoff value to output the final analysis results displayed as forest plots. Given only six matrisome component-encoding genes were examined, multiple hypothesis test correction was unnecessary and thus not performed.

### Statistical analysis for sequencing data

See Bulk gene expression data analysis and scRNA-seq data analysis for details.

## Supplementary figures

## Supplementary tables and videos

**Table S1.** Reference marker genes of cell type and state clusters.

**Table S2.** Curated gene sets of microglia phenotypes.

**Table S3.** DEGs of breast cancer-labeled microglia.

**Table S4.** Microglia GSEA results with BP gene sets.

**Table S5.** Microglia GSEA results without BP gene sets.

**Table S6.** Upregulated DEGs of HER2BC-BrM.

**Supplementary video 1.** 3D animation of immunolabeling-enabled imaging of solvent-cleared organs (iDISCO) showing H2030-BrM metastases, corresponding to Figure S1B (first row).

**Supplementary video 2.** 3D animation of immunolabeling-enabled imaging of solvent-cleared organs (iDISCO) showing MDA231-BrM metastases, corresponding to Figure S1B (second row).

**Supplementary video 3.** 3D animation of immunolabeling-enabled imaging of solvent-cleared organs (iDISCO) showing HCC1954-BrM metastases, corresponding to Figure S1B (third row).

---

**Figure S1.**
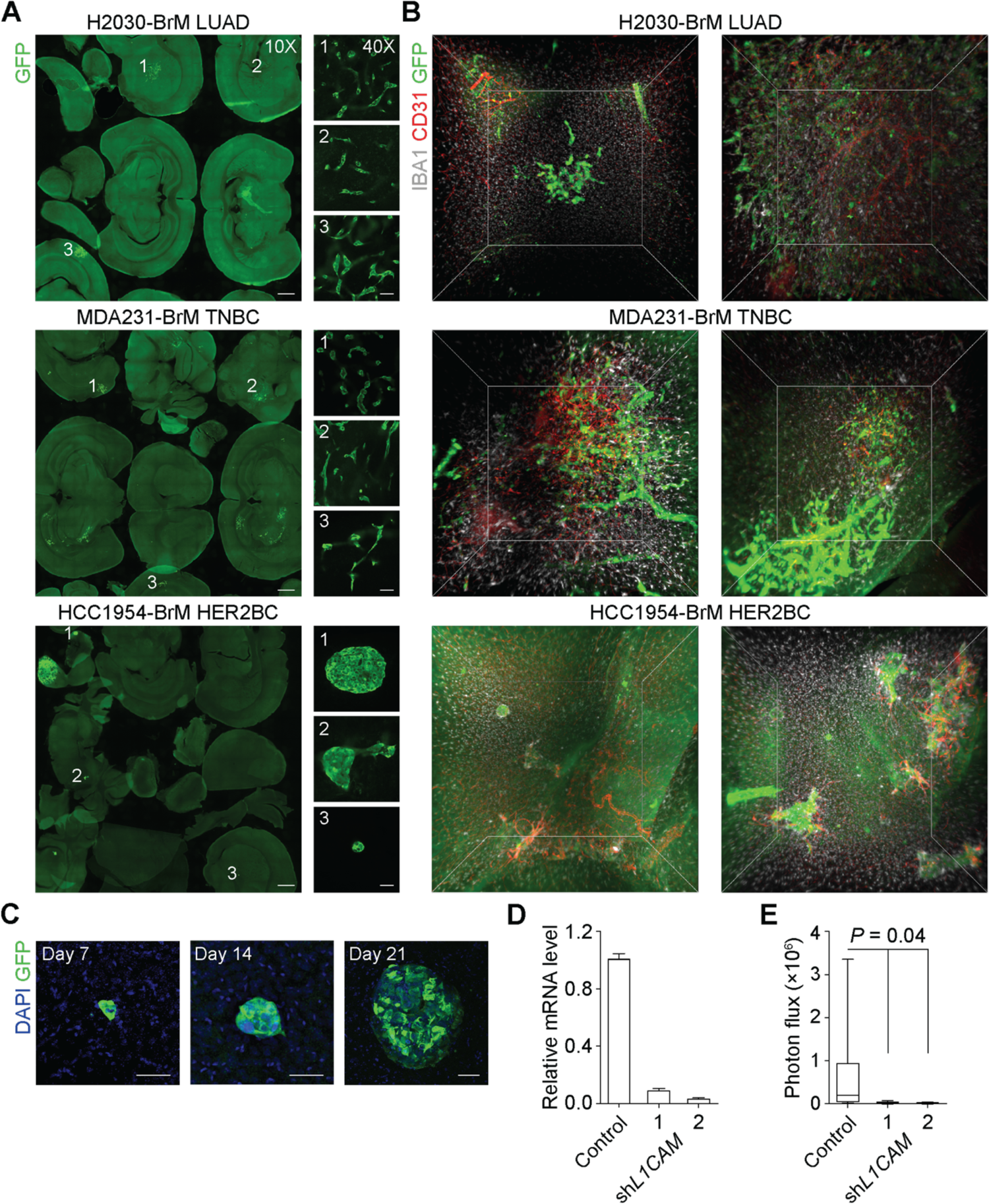
Brain colonization patterns and stromal interface examined by multiple imaging methods and impact of L1CAM on the survival of extravasated HER2BC cells. (A) Representative IF staining of free-floating tissue sections (80 µm) of athymic mouse brains harboring metastases formed by H2030-BrM LUAD, HCC1954-BrM HER2BC, and MDA231-BrM TNBC cells 3-4 weeks post-intracardiac inoculation. Scale bars, (left panels) 1 mm and (right panels) 50 µm in images obtained by 10X and 40X objectives, respectively. The same lesions imaged by both 10X and 40X objectives are indicated by matching numbers, with three example lesions shown for each model. (B) Representative 3D still images of immunolabeling-enabled imaging of solvent-cleared organs (iDISCO) showing cubic brain tissue regions (1 mm^3^) harboring metastases 3-4 weeks post-intracardiac inoculation. Images were obtained by cleared-tissue lightsheet microscopy (Luxendo MuVi SPIM) imaging of whole-mount immunolabeling of a brain hemisphere. Corresponding animation of these still images is shown in Supplementary videos 1-3. (C) Representative IF staining of spheroidal HCC1954-BrM colonies formed at different times after intracardiac inoculation. Scale bars, 50 µm. (D) Relative mRNA levels of *L1CAM* in HCC1954-BrM cells expressing control vector or the indicated 2 shRNAs, measured by qRT-PCR. Mean ± SEM. (E) Effect of *L1CAM* shRNA knockdown (KD) in HCC1954-BrM cells on brain colonization, measured by *ex vivo* BLI of the brain (top panel, n = 6-7 mice/group, 4 weeks post-intracardiac inoculation).

**Figure S2.**
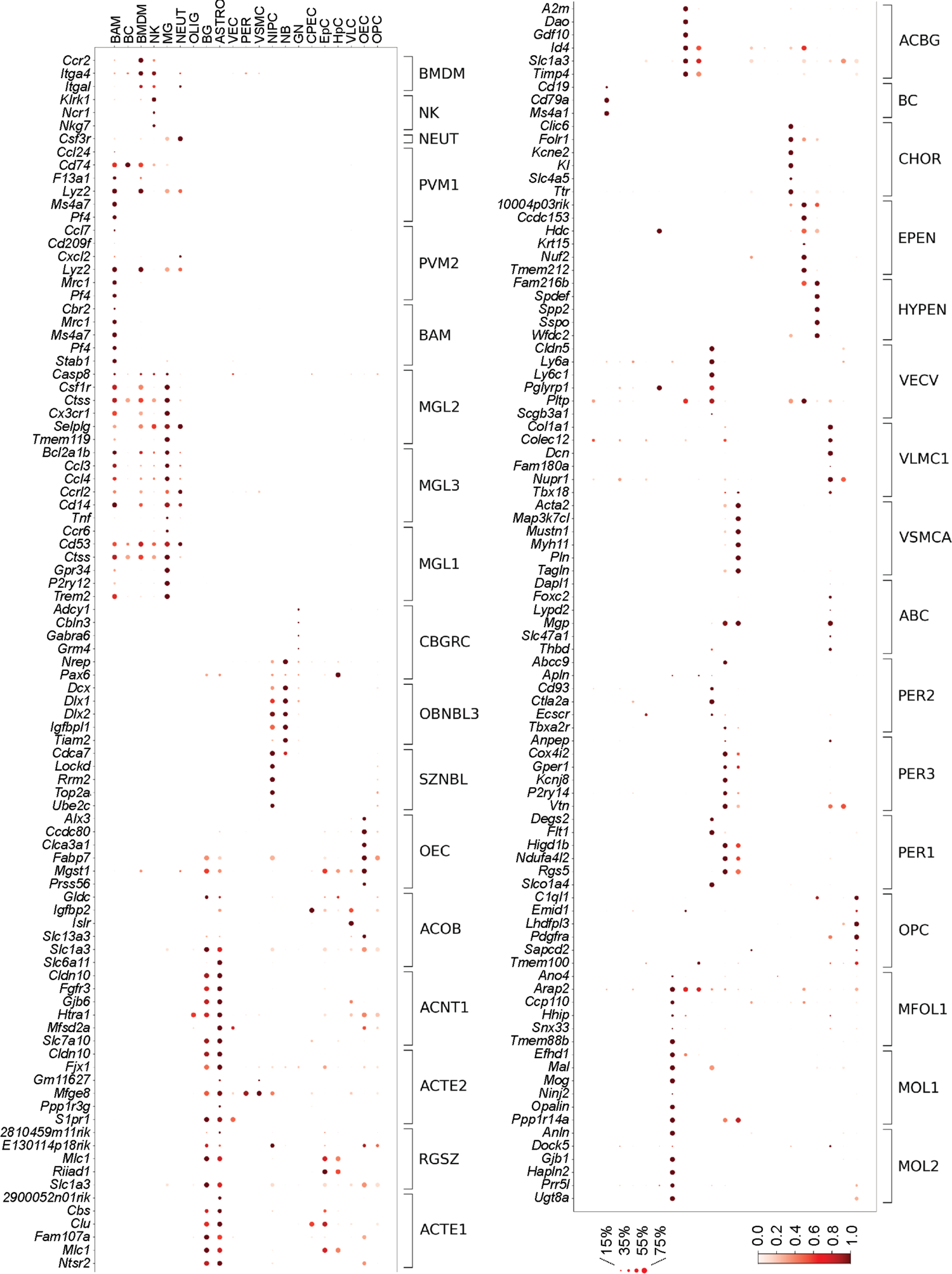
Expression of cell type marker genes in non-cancer cell populations from scRNA-seq profiling. Dot plots showing the expression of marker genes (rows) in each population of non-cancer cells (columns), computed on cells from both scRNA-seq experiment 1 and 2. The size of dots represents the fraction of cells with non-zero expression in each population; and saturation of color indicates the population-averaged log-transformed normalized UMI (unique molecular identifier) counts for each gene standardized to between 0 and 1. The cell populations are shown with identical abbreviated terms in Figure 2D, and in the same order with Figure 2E clustermap and 2F heatmaps (rows). Marker genes were obtained as described in Figure 2.

**Figure S3.**
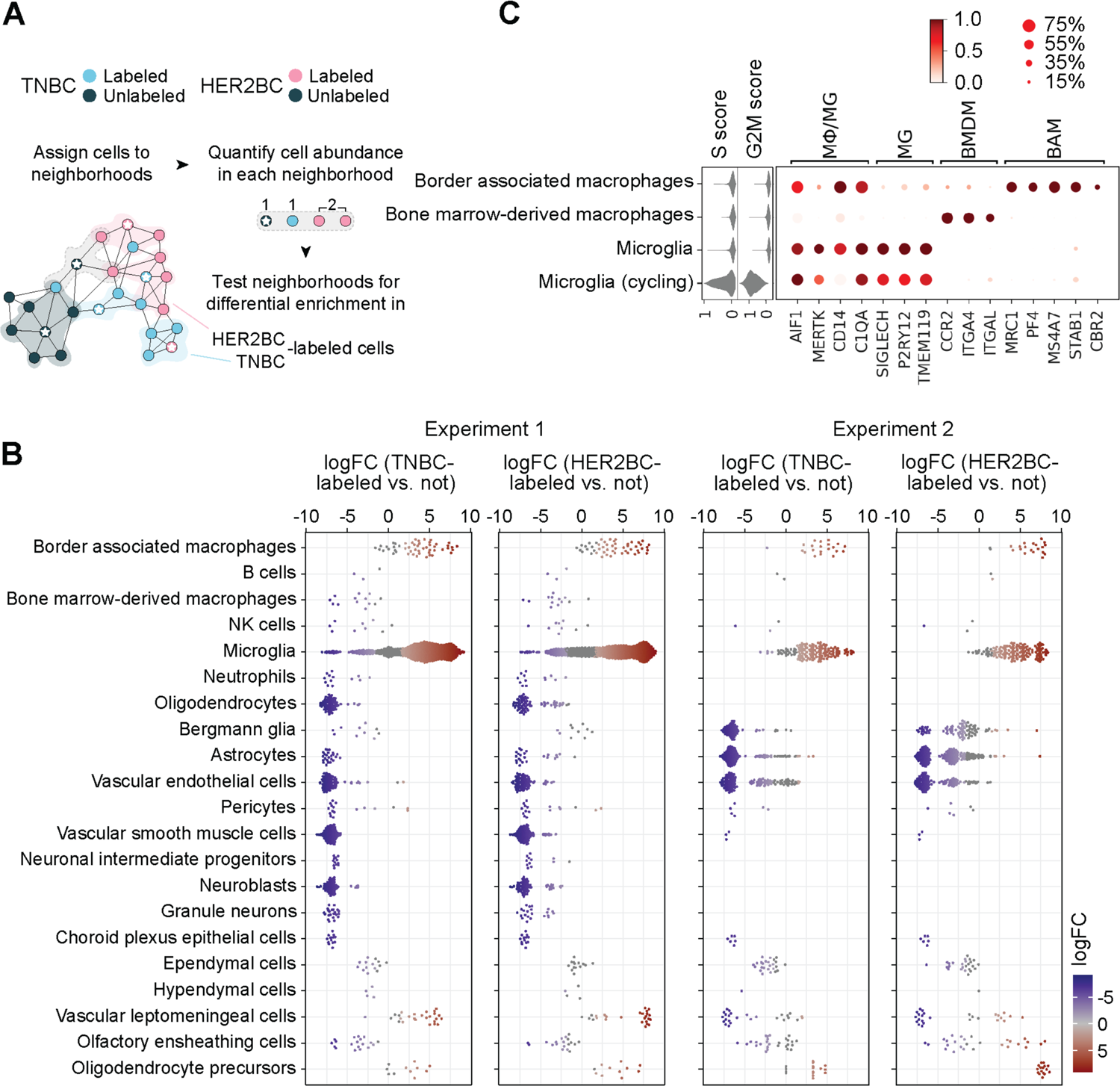
Microglia populations in the TME of MDA231 TNBC and HCC1954 HER2BC brain metastases. (A) Schematic of the statistical framework for quantifying and testing the differential abundance in cells from different sources (adapted from Ref.^47^). Cells (dots) are assigned to partially overlapping phenotypic neighborhoods (encircled by varied background colors, indicative of whether and which source of cells a neighborhood is enriched in) on a *k*-nearest neighbor (*k-*NN) graph. The neighborhoods are defined on index cells (indicated by white stars), selected using a graph sampling algorithm. The counts of cells in the light gray neighborhood (not enriched in cells from any particular source) are shown as an example. (B) Differential abundance (measured by log fold change, logFC, x axis) of TNBC-labeled (left panels) and HER2BC-labeled (right panels) cells in reference to unlabeled cells among all phenotypic neighborhoods, stratified by the types of neighborhood index cells. Results of experiment 1 and 2 are presented in parallel. Phenotypic neighborhoods with significant differential abundance (BH-adjusted P values ≤ 0.1) are colored to indicate corresponding logFC (x axis) values. (C) Expression of canonical cell type markers (shown by dot plot, right panel) and S phase and G2/M phase scores (violin plot, left panel) in the indicated subsets of macrophages. MΦ, macrophages. MG, microglia computationally identified to be in the G1 phase or not cycling. MG (cycling), microglia inferred to be cycling, given high scores of the S and G2/M phases. BAM, border associated macrophages. BMDM, bone marrow-derived macrophages. The size of dots in the dot plot represents the fraction of cells with non-zero expression in each subset of cells; and saturation of color indicates the subset-averaged log-transformed normalized UMI (unique molecular identifier) counts for each gene standardized to between 0 and 1.

**Figure S4.**
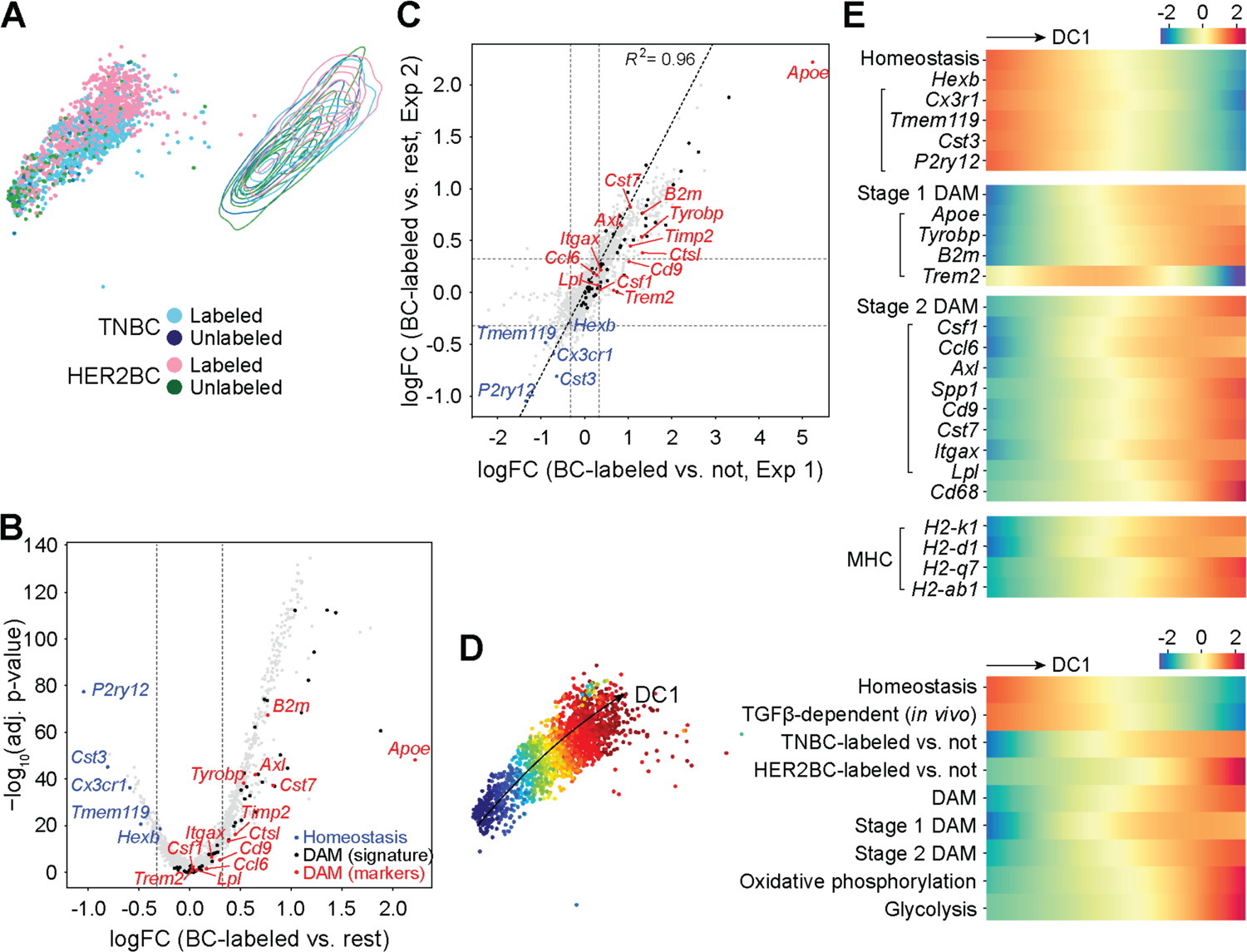
Replication of homeostasis-to-DAM transition in metastasis-associated microglia by an independent scRNA-seq profiling experiment. All results computed on the non-cycling microglia (MG) from experiment 2. (A) (Left panel) UMAP embedding of the 4 indicated sources of altogether 1737 cells, and (right panel) contour plots of each source in the embedding, as in Figure 4A. (B) Volcano plot of the logFC in gene expression against corresponding BH-adjusted P values, comparing the groups of phenotypic neighborhoods concordantly enriched in breast cancer (BC) cell-labeled non-cycling microglia to the rest. 501 (92%) of 546 DEGs detected overlap with the 868 DEGs from experiment 1. (C) Linear regression showing that the logFC values of all 11379 genes with estimable expression coefficients in metastasis-associated non-cycling microglia were well correlated between experiments 1 and 2 (see Table S3 and STAR Methods). x- and y-axis values correspond to the x-axis values in Figure 3C and Figure S4B, respectively. (B, C) As with Figure 3C, logFC thresholds were indicated by gray dashed lines, and sources and lists of indicated gene sets were provided in Table S2. (D) (Left panel) UMAP plot showing first diffusion component (DC1) values computed on all cells by color map; (right panel) heatmap showing fitted trends of indicated neighborhood-level signature scores and labeled cell enrichment (quantified by the local logFC in abundance in reference to unlabeled cells, logFC) along the DC1 values of neighborhood index cells (rows, z-normalized per row across neighborhoods). Marker genes were obtained as detailed in Figure 4F. (E) Heatmap showing fitted trends of neighborhood-level signature scores and gene expression along the DC1 values of neighborhood index cells as shown in the heatmap in (B) (rows, z-normalized per row). Trends of inflammatory cytokine genes *Tnf*, *Ccl4*, *Il1b* expression were not computed given their overall low UMI counts, possibly caused by the addition of transcription and translation inhibitors during the brain tissue harvesting and homogenizing steps in experiment 2.

**Figure S5.**
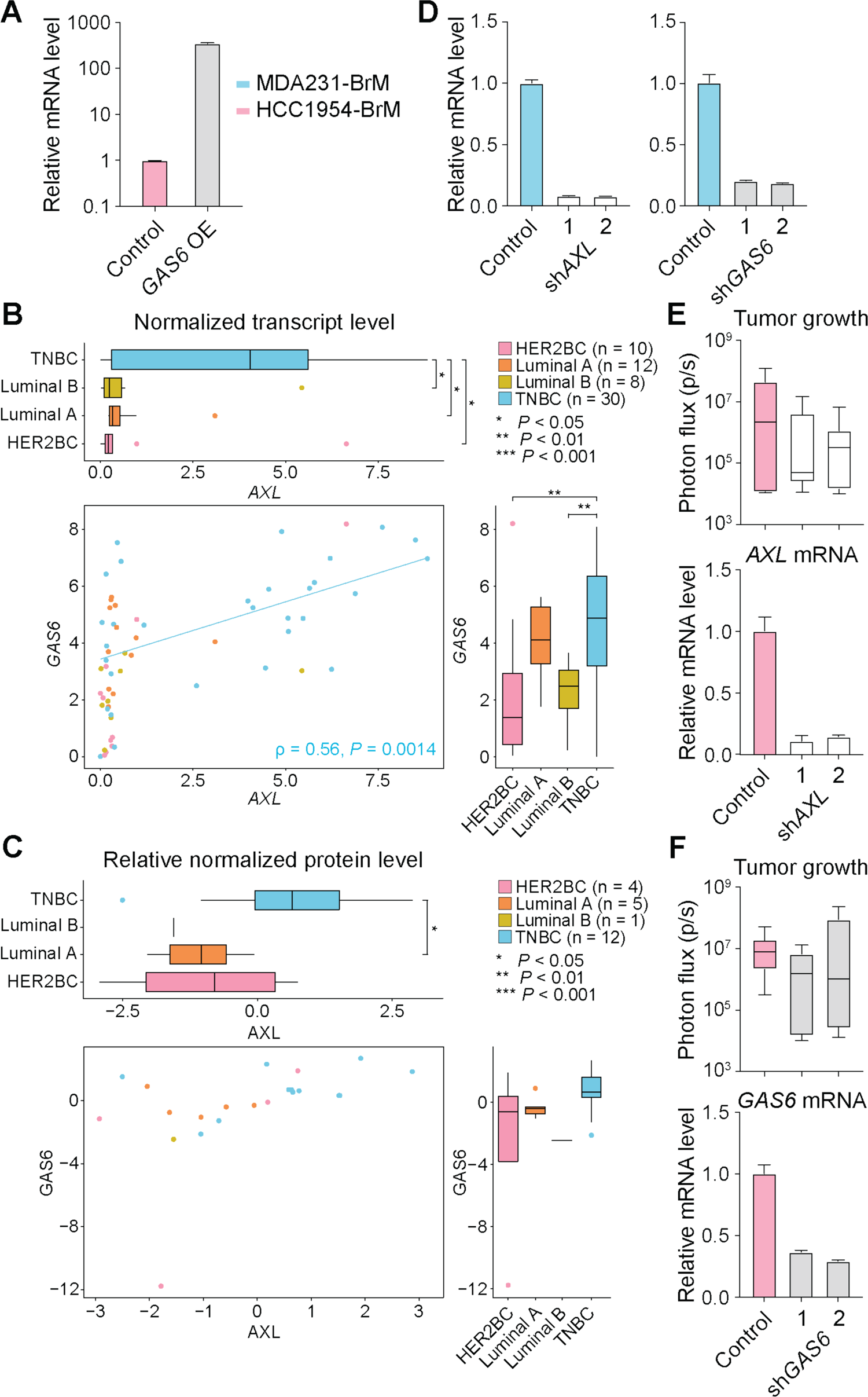
Perturbing GAS6/AXL signaling in MDA231 TNBC and HCC1954 HER2BC brain metastases. (A) Relative mRNA levels of *GAS6* in HCC1954-BrM cells overexpressing (OE) *GAS6* or control vector, measured by qRT-PCR. Mean ± SEM. (B-C) Co-enrichment of GAS6 and AXL in patient TNBC cells. Normalized transcript (B) and relative normalized protein (C) levels of AXL and GAS6 in all available human breast cancer cell lines in DepMap database (depmap.org/portal/). ρ, Spearman’s correlation coefficient. (D) *AXL* (left panel) and *GAS6* (right panel) relative mRNA levels in MDA231-BrM cells expressing control vector or the indicated shRNAs (2 shRNAs per target gene), validated by qRT-PCR. Mean ± SEM. (E-F) Effect of *AXL* (E) and *GAS6* (F) shRNA knockdown (KD) in HCC1954-BrM cells (2 shRNAs per target gene), validated by qRT-PCR (bottom panels, mean ± SEM), on brain colonization measured by *ex vivo* BLI of the brain (top panels, n = 6-7 mice/group, 4 weeks post-intracardiac inoculation).

**Figure S6.**
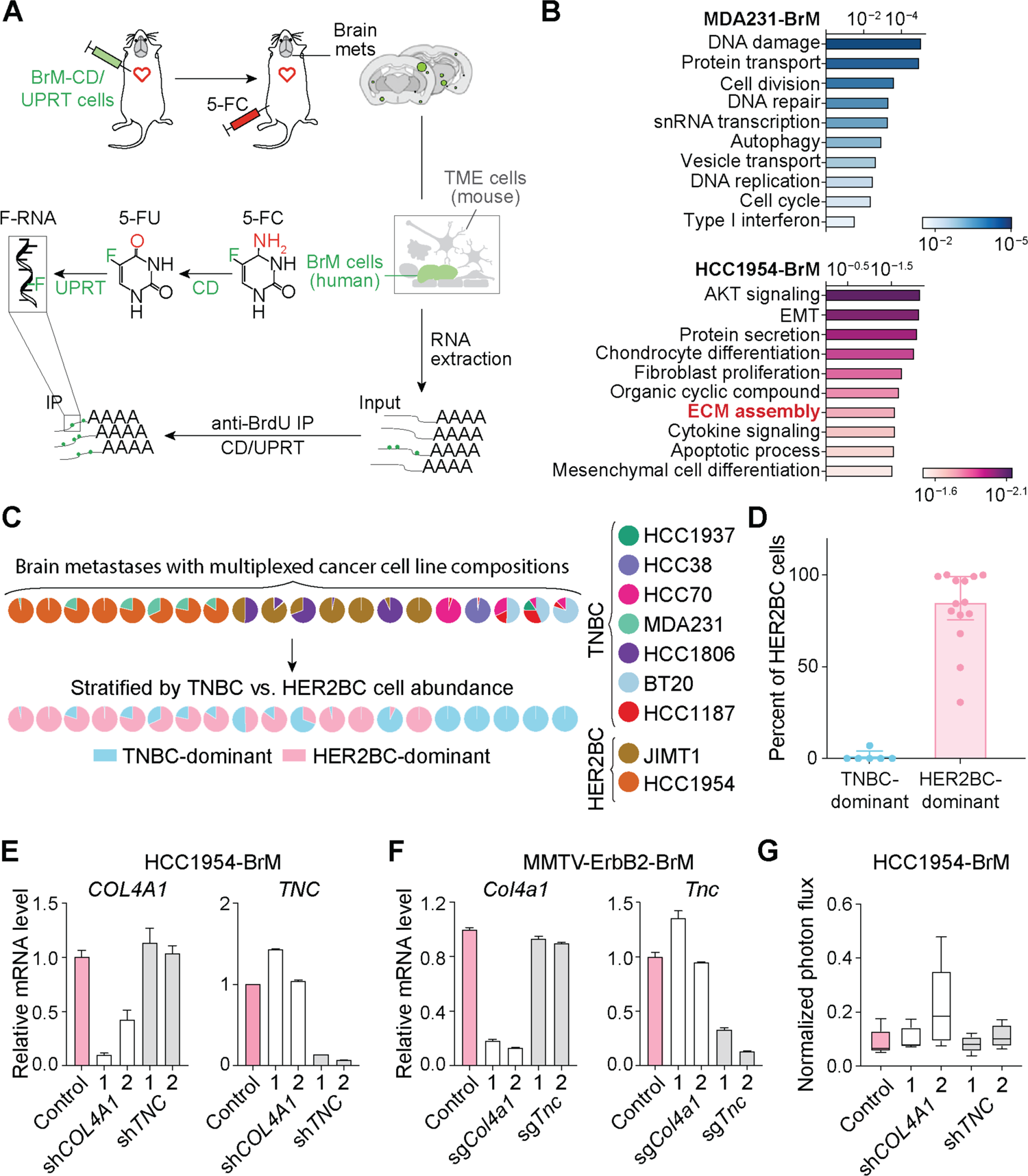
Characterizing and suppressing the expression of matrisome genes in TNBC and HER2BC brain metastases. (A) Schematic illustrating the Flura-seq profiling of *in situ* transcriptome of brain metastases. Athymic mice were inoculated intracardiacally with brain metastatic (BrM) cells engineered to co-express cytosine deaminase (CD) and uracil phosphoribosyl transferase (UPRT). Mice were administered with 5-fluorocytosine (5-FC) (250 mg/kg) by intraperitoneal (IP) injection 12 hours prior to harvesting the brain tissue. 5-FC was converted through a series of steps to 5-fluorouridine triphosphate (F-UTP) specifically in CD/UPRT expressing BrM cells. F-UTP was incorporated into nascent RNA of the cells to yield the 5-FU-tagged RNA (F-RNA) that could be purified by anti-BrdU immunoprecipitation for RNA-seq analysis. Specifically, CD converted 5-FC to 5-fluorouracil (5-FU); and UPRT converted 5-FU to cell membrane-impermeable 5-fluorouridine monophosphate (F-UMP), which was ultimately converted to 5-fluorouridine triphosphate (F-UTP) by endogenous enzymes. (B) Overrepresentation analysis showing the top gene ontology (GO) terms associated with genes respectively upregulated in MDA231 and HCC1954 brain metastases, compared in reference to each other (see STAR Methods). Color shades indicate BH-adjusted P values of normalized enrichment scores. (C) Composition of cells from indicated TNBC and HER2BC cell lines in brain metastases formed by each multiplexed cell line pool, quantified by deep sequencing of the barcodes of cell lines. (D) Percentage of HER2BC cells in multiplexed brain metastasis samples classified to be dominated by TNBC (< 25% HER2BC cells) and HER2BC cells, respectively. Median with interquartile range. (E) *TNC* and *COL4A1* relative mRNA levels in HCC1954-BrM cells expressing control vector or the indicated shRNAs (2 shRNAs per target gene), measured by qRT-PCR. Mean ± SEM. (F) *Tnc* and *Col4a1* relative mRNA levels in MMTV-ErbB2-BrM cells expressing control vector or the indicated CRISPRi sgRNA, measured by qRT-PCR. Mean ± SEM. (G) Effect of *TNC* and *COL4A1* shRNA knockdown (KD) on lung colonization by HCC1954-BrM cells. Lung photon flux was measured 4 weeks post-tail vein inoculation (n = 5 mice/group).

