## Supplementary table 6 for "Distinct tumor architectures for metastatic colonization of the brain"

**Table S6. Genes upregulated in both HCC1954 and MMTV-ErbB2 BrM derivatives, compared to their corresponding parental (Par) cell lines.**

| **Gene** | **Fold change of BrM in reference to Par** | |
| --- | --- | --- |
|  | **HCC1954** | **MMTV-ErbB2** |
| *SEMA7A* | 3.44 | 27.91 |
| *KCNK2* | 2.97 | 24.33 |
| *BDKRB2* | 2.07 | 24.09 |
| *LAMC2* | 5.97 | 7.24 |
| *SPNS3* | 2.50 | 13.70 |
| *APCDD1* | 2.13 | 15.35 |
| *RASGRP1* | 6.48 | 3.50 |
| *EXOC3L4* | 3.30 | 6.35 |
| *NOS3* | 3.01 | 6.39 |
| *S100A7A* | 4.00 | 4.54 |
| *H2AFY2* | 4.47 | 3.94 |
| *IL24* | 2.09 | 8.05 |
| *TNC* | 4.45 | 6.87 |
| *B4GALNT3* | 2.80 | 4.81 |
| *CRIP2* | 2.32 | 5.66 |
| *PADI2* | 2.39 | 5.02 |
| *IRF8* | 2.14 | 5.57 |
| *DUSP5* | 3.13 | 3.58 |
| *GNAZ* | 2.99 | 3.20 |
| *PTPRE* | 3.11 | 2.99 |
| *IRS2* | 4.24 | 2.10 |
| *GYLTL1B* | 2.05 | 4.28 |
| *KLHL38* | 2.61 | 3.33 |
| *COL4A1* | 2.04 | 4.00 |
| *PIPSK1B* | 2.03 | 3.88 |
| *PDZD4* | 2.85 | 2.70 |
| *ERRFI1* | 2.61 | 2.34 |
| *CST6* | 2.11 | 2.79 |
| *SDPR* | 2.01 | 2.71 |
| *EEPD1* | 2.16 | 2.45 |
| *CSGALNACT1* | 2.18 | 2.38 |
| *HS3ST1* | 2.15 | 2.34 |
